# Unstable anthocyanin pigmentation in *Streptocarpus* sect. *Saintpaulia* (African violet) is due to transcriptional selectivity of a single MYB gene

**DOI:** 10.1101/2025.01.10.632376

**Authors:** Daichi Kurata, Tomohisa Tsuzaki, Fumi Tatsuzawa, Kenta Shirasawa, Hideki Hirakawa, Munetaka Hosokawa

**Affiliations:** Graduate School of Agriculture, Kindai University, 3327-204, Nakamachi, Nara, Nara 631-8505, Japan; Faculty of Agriculture, Iwate University, 3-18-8, Ueda, Morioka, Iwate 606-8550, Japan; Kazusa DNA Research Institute, 2-6-7, Kazusa-Kamatari, Kisarazu, Chiba 292-0818, Japan; Agricultural Technology and Innovation Research Institute Kindai University (ATIRI), 3327-204, Hanamachi, Nara, Nara 631-8505, Japan

**Keywords:** Epigenetics, Periclinal chimera, Alternative Promoter Usage, Anthocyanin

## Abstract

- *Saintpaulia* (African violet) pigmentation is notoriously unstable, particularly following passage through tissue culture, but the instability mechanism is unknown. White-striped petals were thought to be due to epigenetics, but the genes involved are unknown. We confirmed that white stripes result from epigenetic regulation based on the flower color traits of plants obtained from tissue culture.
- Gene expression in several plant lines, anthocyanin quantification, bisulfite sequencing, and methylation analyses were used to demonstrate the presence of a single MYB gene responsible for pigment variation. We identified *SiMYB2* as the cause of variations in tissue color patterning, and that two RNAs were generated from *SiMYB2*. *SiMYB2-Long* was expressed in colored tissues, while *SiMYB2-Short* was expressed only in non-colored tissues. Functional analyses revealed that SiMYB2-Long is an anthocyanin biosynthesis activator and SiMYB2-Short is non-functional. Exon 3 of *SiMYB2* was generated by the insertion of a transposon-like sequence. A mutant lacking the element was obtained from cultivars with non-colored tissues. Anthocyanin content and *SiMYB2-Long* expression in the mutant were greatly increased compared to wild type.
- Our results suggest that the white-striped petals of *Saintpaulia* are not formed by periclinal chimeras but thorough the transcriptional selectivity of epigenetically regulated *SiMYB2*.

## Introduction

African violet (*Streptocarpus* sect. *Saintpaulia*) is an ornamental plant native to Kenya and Tanzania (Möller & Cronk, 1997). *Saintpaulia* accumulates anthocyanin in its petals, resulting in diverse color patterns such as stripe, picotee, splash, center eye, and spots. Notably, periclinal chimeras with bicolor striped petals are valuable due to the difficulties associated with their propagation (Eyerdom, 1981). *Saintpaulia* is typically propagated via adventitious shoots from tissue culture or leaf cuttings. However, in many cases adventitious shoots originate from multiple epidermal cells, leading to monochromatic petals rather than stripes when pericarpal chimaeras are propagated through adventitious buds (Broertjes & Keen, 1980; Ohki, 1994; Nabeshima *et al*., 2017; Yang *et al*., 2017). Thus, propagation of polychromatic periclinal chimera patterns through adventitious shoots is not possible.

Although some *Saintpaulia* cultivars have striped petals similar to those observed in periclinal chimeras, the plants regenerated from them often exhibit varying color patterns. The petals of these cultivars exhibit a pinwheel-like pattern with a central-white region surrounded by a pigmented marginal region. Many white-striped cultivars have been bred and have become a popular *Saintpaulia* pigmentation pattern. These white-striped cultivars were thought to be periclinal chimeras (Lineberger & Druckenbrod, 1985; Kazemian *et al*., 2019), but plants regenerated from these cultivars sometimes have white-striped petals, in addition to monochromatic petals, which makes it difficult to conclude that the white stripes are due to periclinal chimeras. Regenerated plants with white petals have unstable pigmentation and may also produce petals that are either white-striped or of a uniform color. Therefore, the mechanism of pattern formation in white-striped petals may be attributed to the instability of gene expression as a result of changes in epigenetic regulation (Yang *et al*., 2017).

Ong-Abdullah *et al*. (2015) identified a gene (*EgDeficiens1*) that controls a trait that results in malformed fruit produced via tissue culture in African oil palm (*Elaeis guineensis*). In addition, Tang et al. (2022) identified a gene (*CmMYB6*) responsible for floral color variation emerging from lateral buds in Chrysanthemum (*Chrysanthemum morifolium*). Because both of these studies successfully used cytosine methylation analysis to identify the epigenetic target, genes with different methylation levels among different traits are likely candidates as the causal genes. In this study we have investigated the long-enigmatic phenomenon of pigment accumulation and pattern formation in *Saintpaulia*, based on the hypothesis that they result from epigenetic mutations. Using an integrated approach combining gene expression analysis and epigenomic profiling, we aimed to identify the gene or genes underlying this instability.

## Materials and methods

### Plant materials and tissue culture

*Saintpaulia* cultivar ‘Kilauea’ has white-striped petals, and we obtained plants with pink petals via tissue culture of this cultivar (Fig. 1a, b). Propagated plants with pink petals were labeled as ‘KP’. Leaves collected from KP plants were surface sterilized with 1% sodium hypochlorite supplemented with 10 μl Tween 20 for 15 min. The leaves were then rinsed with double-distilled water and cut into 1–2 cm^2^ pieces using a razor. To obtain adventitious shoots, the pieces were cultured on full-strength MS medium (Murashige & Skoog, 1962) containing 2% sucrose (g/v), 0.3% gellan gum (g/v), 0.1 ppm 1-naphthalene acetic acid (NAA), and 0.5 ppm 6-benzyladenine (BA) for four weeks. Regenerated plants with white petals were labeled ‘KW’ (Fig. 1c), and those with white-striped petals were labeled ‘KWS’ (Fig. 1d). The cultivar ‘Stephanie’ (ST) was used as a comparator line. Stephanie plants emerging from leaf cuttings with white petals (SW), and with white-striped petals (SWS) were labeled accordingly (Fig. 1e, f, g). Cultivars ‘Akira’ (AK), ‘Tomoko’ (TK), ‘Georgia’ (GG), ‘Barbara’ (BB), ‘Manitoba’ (MT), ‘Taro’ (TR), ‘Saturn’ (SA), ‘Shimai’ (SH), and mutants of SH (m-SH) were also included for anthocyanin assays and gene analyses (Fig. 7d, e, Fig. S8a-i). All plants were grown at 21±2 °C, 14 h light/10 h dark, under fluorescent light (BioLux, 20W, HotaluX, approximately 50 μmol cm^-2^ s^-1^). The plants were planted in 220 ml plastic pots filled with Metro-Mix 350J soil (HYPONeX, Osaka, Japan), and fertilizer (HYPONeX N-P-K = 6-10-5, HYPONeX) was applied once every two weeks.

**Fig. 1.**
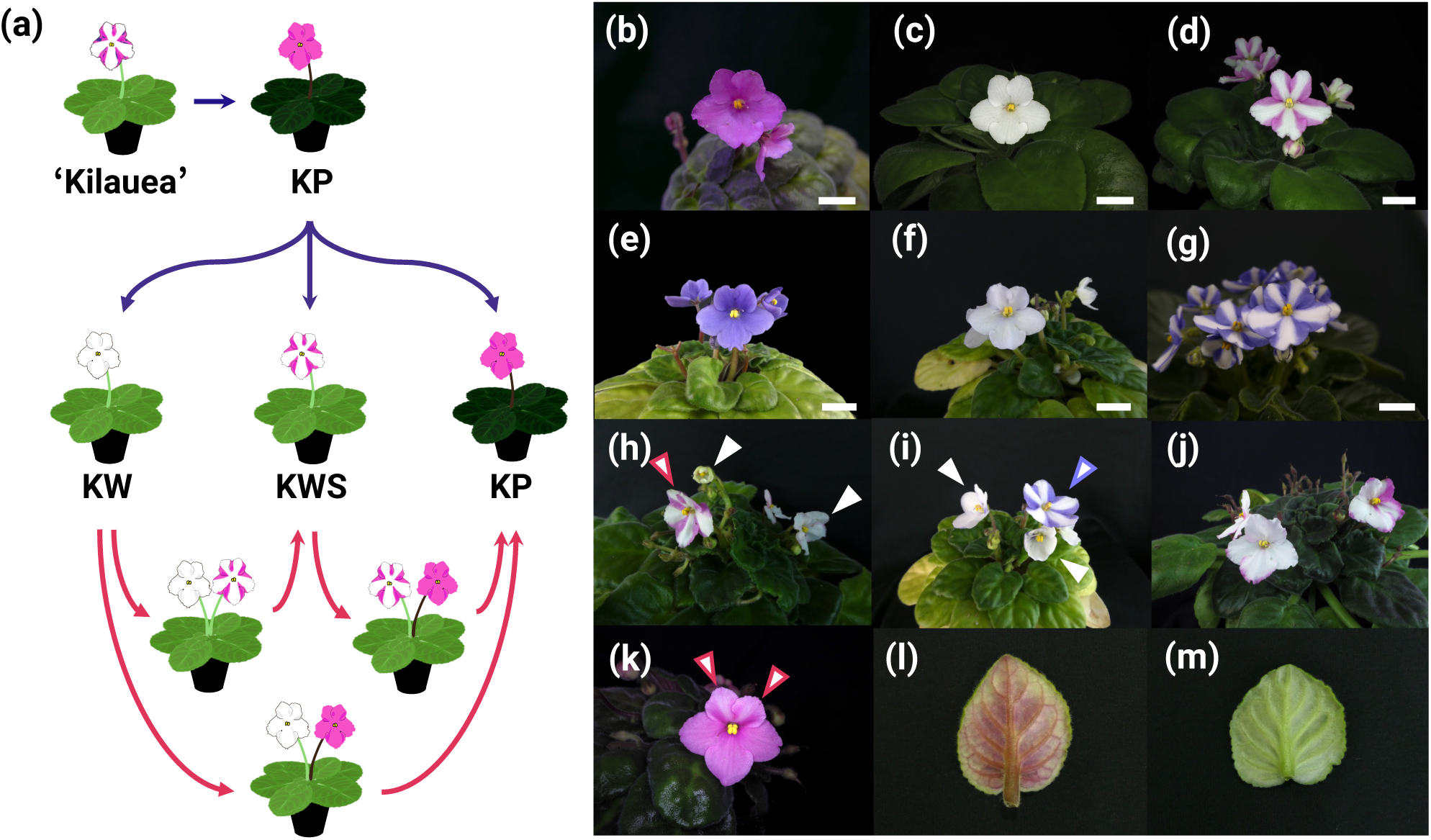
(a) Variation in petals and leaf color of *Saintpaulia* ‘Kilauea’ that occurs via tissue culture (blue arrow), and changes that occur during cultivation (red arrow). (b-g) Flowers of *Saintpaulia*. White bars indicate 1.5 cm scale. (b) KP. (c) KW. (d) KWS. (e) ST, (f) SW, (g) SWS. (h, i) White and white striped flowers are present on a single plant simultaneously. A white arrowhead indicates a white flower and an arrowhead with colored margins indicates a white-striped flower. (j) Randomly pigmented KW petals. (k) Entirely pigmented KWS petals. (l) Leaf of KP. (m) Leaf of KW.

KW leaves were cultured on MS medium as above for four weeks to obtain adventitious shoots. If KWS plants were chimeric with pink pigmentation in the L1 layer and white (*i.e.* no pigmentation) in the L2 layer and below, all shoots originating from the L2 layer and below would be expected to display a white monochromatic phenotype. Tissues, including the epidermal cell layer (L1) of KWS petioles were excised using a razor and forceps, leaving only the subepidermal cell layer(s) (L2 or below). The resulting explants were cultured in MS medium for three months to obtain adventitious shoots. All regenerated plants were grown as described above until they reached the flowering stage.

### Flavonoid analysis

To identify which stage of the anthocyanin biosynthetic pathway was interrupted in KW petals, the anthocyanin and flavone compositions of KP and KW petals were measured by high-performance liquid chromatography (HPLC). Petals of KP and KW were collected 2–3 days after flower opening and frozen at -80 °C. The frozen petals were quickly crushed using a mortar and pestle at room temperature to avoid volume increases due absorption of additional moisture from the air. The powdered petals were then extracted with ten-fold weight of 50% acetic acid-methanol (acetic acid: methanol: distilled water = 1:4:5, v/v/v), and filtered through a 0.45 μm syringe filter (SLNY1345NB, Nippon Genetics, Tokyo, Japan). The filtered solution was then subjected to HPLC for qualitative and quantitative determination of flavonoids. Analytical HPLC was performed using a C18 (4.6 × 250 mm) column (Waters, Milford, Massachusetts, USA) at 40 °C with a flow rate of 1 mL/min, monitored by a photodiode array detector on an LC 10A system (Shimadzu Corporation, Kyoto, Japan). Linear gradient elution was performed using an acetic acid-methanol mobile phase, from 20 to 85% solvent B (1.5% H_3_PO_4_, 20% HOAc, 25% MeCN in H_2_O) in solvent A (1.5% H_3_PO_4_ in H_2_O) over 40 minutes, followed by a five-minute re-equilibration at 20% solvent B for flavonoids (530 and 350 nm).

Absorbance at 530 nm (Abs_530_) was measured using a Shimadzu UV-1800 spectrophotometer (Shimadzu, Japan) for the anthocyanin extracts derived from the petals of KP, KW, ST, and SW. A calibration curve was constructed from a five-fold dilution series of cyanidine-3-glucoside. Petal anthocyanin content was quantified as cyanidine-3-glucoside equivalents using the calibration curve. The 5th to 7th expanded leaves of KP, KW, SH and m-SH were collected to measure their anthocyanin content, then extracted and measured as described above.

### qRT-PCR and RT-PCR

The expression of anthocyanin biosynthesis-related genes was investigated by quantitative RT-PCR (qRT-PCR). Petal development was divided into four stages, designated as S1 to S4 (Fig. S1). Petals covered by sepals and showing no coloration were designated as S1, the beginning of coloration as S2, the development of deeper coloration and just before flowering as S3, and fully colored petals around the time of flower opening as S4. Petal buds of KP, KW, ST, and SW buds of S2 and the youngest leaves of KP and KW were collected, frozen in liquid nitrogen, and total RNA was extracted using Sepasol^®^-RNA I Super G (Nacalai Tesque, Kyoto, Japan). First-strand cDNA was synthesized from total RNA using oligo dT primers and ReverTra Ace^®^ (TOYOBO, Osaka, Japan). Reverse transcription reactions were performed at 42 °C for 30 min and 99 °C for 5 min. qRT-PCR was performed using cDNA and SYBR THUNDER BIRD^®^ qPCR Mix (TOYOBO) by CFX ConnectTM Real-Time PCR Detection System (Bio-Rad, Hercules, CA, USA). Reaction conditions were 95 °C 10 sec and 60 °C 30 sec for 40 cycles. Primers specific for anthocyanin biosynthesis-related genes were designed using Primer 3 Plus (Untergasser *et al*., 2012) based on BLAST and annotation information of predicted gene sequences (Table S1). A calibration curve with an R^2^ ≥ 0.98 was constructed, and relative quantification was performed. *SiActin* (Sio_r2.0_p0205.1.g00590.1) was used as the housekeeping gene for normalization.

Petals of KWS and SWS were collected on the day of flower opening. Pigmented and non-pigmented regions of the petals at S4 were separated with a razor and immediately frozen in liquid nitrogen. RNA was then extracted and qRT-PCR was performed as described above.

RT-PCR was performed to detect the expression of *SiMYB1* in petals and leaves. Petals (at S2) and leaf RNAs were reverse transcribed as described above. PCR reactions were performed using KOD One^®^ PCR Master Mix (TOYOBO). Reactions were performed at 98°C for 10 sec, 60°C for 5 sec, and 68°C for 1 sec for 35 cycles and amplified to a plateau. Primers were the same as those used for qRT-PCR (Table S1). *SiActin* was used as the internal control.

### RNA-seq analysis

RNA-seq was performed to analyze the transcriptome of KP and KW. Petals were collected from KP and KW buds (S2) and frozen in liquid nitrogen. Total RNA was extracted using the RNeasy Plant Mini Kit (Qiagen, Venlo, The Netherlands) according to the manufacturer’s instructions. RNA-seq library preparation and sequencing were performed by the Rhelixa genomic and epigenomic research services company (Tokyo, Japan). RNA-seq libraries were prepared using the NEBNext^®^ Poly(A) mRNA Magnetic Isolation Module and the NEBNext^®^ Ultra^TM^ II Directional RNA Library Prep Kit (New England Biolabs, Ipswich, MA, USA). 150 bp paired-end sequencing was performed on an Illumina NovaSeq 6000 (Illumina, San Diego, CA, USA), yielding 4 Gb reads. Adaptor sequences and low-quality reads were removed computationally using Trimmomatic v0.39 (Bolger *et al*., 2014).

The *Saintpaulia* SIO_r2.0 genome data (derived from the KP genome, with 37,682 predicted genes. BioProject PRJDB15710) was used as the reference sequence (Kurata et al., 2024). Mapping to SIO_r2.0 was done using HISAT2 v2.2.1 (Kim *et al*., 2019). Read count data and transcripts per million (TPM) values were calculated by Stringtie v2.2.1 (Pertea *et al*., 2015). Differentially expressed genes (DEG) analysis using TCC package (Sun *et al*., 2013) and edgeR v4.0.3 (Robinson *et al*., 2010) was performed using read count data with the TMM method (FDR = 0.05). Mapping results obtained by RNA-seq analysis were visualized by pyGenomeTracks v3.8 (Lopez-Delisle *et al*., 2021). De novo assembly of transcript sequences was performed by Trinity v2.14.0 (Grabherr *et al*., 2011) using RNA-seq data.

### Phylogenetic tree construction

Molecular phylogenetic trees were constructed for MYB and bHLH transcription factors (TFs), which are important for regulating the expression of anthocyanin biosynthesis genes (ABGs). Amino acid sequences were filtered using PREQUAL v1.02 (Whelan *et al*., 2018), and aligned using MAFFT v.7.490 (Katoh & Standley, 2013) with default settings. Multiple aligned sequences were trimmed using trimAl v1.4.1 (Capella-Gutierrez *et al*., 2009), and used to generate a molecular phylogenetic tree by IQ-TREE2 v2.2.0.3 (Minh *et al*., 2020). Internal branch robustness was determined with Ultrafast bootstrapping (UF-boot) and SH-aLRT 1,000x each, and branches with ≥ 95% and ≥ 80% confidence levels were considered robust (Minh *et al*., 2013). The classification of MYB and bHLH TFs was based on Stracke et al. (2001) and Heim (2003), respectively. The accession IDs of the amino acid sequences are listed in Table S2.

### Whole-genome bisulfite sequencing (WGBS)

Unstable pigmentation in petals and leaves may result from changes in the epigenomic state of genes in those organs. Therefore, WGBS was used to identify differences in the methylated cytosine status of anthocyanin biosynthesis-related genes. The first through fourth expanded leaves were collected from KP and KW plants. Leaves were flash-frozen in liquid nitrogen and DNA was extracted using QIAGEN Genomic-tip 100/G (Qiagen). The DNA was treated with bisulfite using an EZ DNA Methylation Gold Kit (Zymo Research, CA, USA). WGBS library preparation and sequencing were performed by Rhelixa, Inc. WGBS libraries were prepared from bisulfite-treated DNA using an Accel-NGS^®^ Methyl-Seq DNA Library Kit (Swift Biosciences, MI, USA), and 150 bp paired-end sequences were generated on an Illumina NovaSeq 6000 or NovaSeq X plus (Illumina), yielding 30 Gb reads. Adaptor sequences and low-quality reads were removed using Trim Galore v0.6.7 (https://github.com/FelixKrueger/TrimGalore). The *Saintpaulia* SIO_r2.0 genome was used as the reference sequence (Kurata et al., 2024). WGBS sequences were mapped to SIO_r2.0 using BWAmeth v0.2.7 (arxiv.org/abs/1401.1129). Then, CG, CHG, and CHH methylation contexts were extracted using MethylDackel v0.6.1 (https://github.com/dpryan79/MethylDackel). Differentially methylated regions (DMRs) in the Sio_r2.0_p0007.1 contig were obtained using methylKit v0.99.2 (Akalin *et al*., 2012). Methylation levels were visualized by pyGenomeTracks v3.8 (Lopez-Delisle *et al*., 2021).

### McrBC-PCR/qPCR

*SiMYB2* methylation levels were compared between pigmented and non-pigmented tissues using McrBC, a methylation-dependent restriction endonuclease. DNA was extracted from the first through fourth expanded leaves of KP and KW using NucleoBond^®^ HMW DNA (Takara, Shiga, Japan) and from the petals of KP and KW using a FavorPrep^TM^ Plant Genomic DNA Extraction Mini Kit (Favorgen, Ping Tung, Taiwan), and digested according to the method described by Jaligot *et al*. (2014). Briefly, 100 ng of DNA was digested with 3 U McrBC (Takara) at 37 °C overnight. The DNA was then incubated at 65 °C for 20 min to stop the reaction and used as template for PCR. Genomic DNA without McrBC treatment was used as a control. The primers used are listed in Table S1. PCR and qPCR reactions were performed using KOD One^®^ PCR Master Mix (TOYOBO) and SYBR THUNDER BIRD^®^ qPCR Mix (TOYOBO), respectively.

### Cloning of *SiMYB2-Long*

First-strand cDNA was synthesized from RNA extracted from KP petals using oligo dT primers and ReverTra Ace^®^ enzyme (TOYOBO). The full-length sequence of *SiMYB2-Long* was amplified using cDNA as a template with Blend Taq^®^ enzyme (TOYOBO). Primer sequences are listed in Table S1. Amplicons were ligated to pTAC-1 using the DynaExpress TA PCR Cloning Kit (BioDynamics Laboratory, Tokyo, Japan), transformed into *Escherichia coli* DH5α competent cells, and colonies carrying the target fragments were selected. Plasmids were extracted from the selected colonies using FastGene^TM^ PlasmidMini (NIPPON Genetics, Tokyo, Japan), and the DNA sequences were confirmed with Sanger sequencing by AZENTA (Tokyo, Japan).

### CAGE-seq analysis

Cap Analysis of Gene Expression (CAGE)-seq was performed to determine the transcription start site (TSS) of *SiMYB2*. Three petal buds were collected at S1 from KP and KW plant lines and frozen in liquid nitrogen. Total RNA was extracted using Sepasol^®^-RNA I Super G (Nacalai Tesque). CAGE library preparation, sequencing, and mapping were performed by DNAFORM (Kanagawa, Japan). cDNA was synthesized from total RNA using random primers. Ribose diols in the 5’ cap structures of RNA molecules were oxidized and biotinylated. The biotinylated RNA/cDNA complexes were then isolated using streptavidin beads (cap-trapping method). Following RNA digestion with RNaseONE/H and adaptor ligation at both cDNA ends, double-stranded cDNA libraries (CAGE libraries) were generated and sequenced using 75 nt single-end reads (NextSeq 500, Illumina). The resulting reads were mapped to the *Saintpaulia* SIO_r2.0 genome using STAR v2.7.9a (Dobin *et al*., 2013).

### Agroinfiltration assays

*Nicotiana tabacum* was grown for transient assay by sowing seeds in Kumiai Nippi Gardening Soil No. 1 (Nihon Hiryo, Tokyo, Japan) and germinating them under 20 °C, 14 h light/10 h dark conditions. The germinated seedlings were transferred to 220 ml plastic pots filled with the same soil and grown under the same conditions for 4–5 weeks.

Four distinct vectors (*SiMYB1*, *SiMYB2-Long*, *SiMYB2-Short*, *SibHLH1*) were designed for transient assays. Vectors were constructed by artificial synthesis using VectorBuilder (VectorBuilder Japan, Kanagawa, Japan) or the in-fusion method. Each vector was designed to have the CDS of each TF driven by the CaMV35S promoter and the kanamycin marker gene NPT II. The VectorBuilder ID of each vector is listed: *SiMYB1* ID; VB210520-1064sfu, *SiMYB2-Long* ID; VB210520-1068fhv, *SibHLH1* ID; VB210520-1053zmv.

The pBI121 vector was digested with *Bam*HI (Takara) and *Sac*I (Takara) to remove the *GUS* coding region. The CDS fragment of *SiMYB2-Short* was amplified using cDNA obtained by reverse transcription of petal RNA. CDS fragments were then amplified using primers for the in-fusion system (Table S2) (In-Fusion^®^ HD Cloning Kit, Takara Bio USA, CA, USA) by combining pBI121 digested fragments and *SiMYB2-Short* amplicons. In-fusion cloning was performed according to the manufacturer’s protocol.

Vector constructs were introduced into Agrobacterium (*Rhizobium radiobactor*) strain EHA105. Transformed Agrobacterium were grown on LB medium (0.5% Yeast Extract, 1% Tryptone, 1% NaCl, 100 ppm Kanamycin, 100 ppm Rifampicin) for two days, and a single colony was selected for the transient assay. Vectors carrying *Green fluorescence protein* (*GFP*) were introduced into Agrobacterium and used as negative controls.

Agroinfiltration was performed with a slight modification of the method of Outchkourov et al. (2014). Transformed Agrobacterium were suspended in liquid YEP medium amended with 50 ppm kanamycin and incubated overnight at 28 °C with shaking. The culture medium was centrifuged (3,500g, 10 min), pellets were resuspended in infiltration buffer [10 mM MES-NaOH (pH = 5.7), 10 mM MgCl_2_, 150 μM acetosyringone], and adjusted to OD_600_ = 0.4–0.6. The mixture was incubated for 2–3 hours at room temperature. The mixture was then injected into 4–5-week-old tobacco leaves. Leaves were photographed, collected and stored frozen at -80 °C three days post-infiltration. The experiment included three biological replicates of two leaves per tobacco plant, and the experiment was repeated in triplicate. Anthocyanin content was then quantified according to the procedure described above.

Leaf discs were collected from the injection site using a biopsy punch (BP-60F, 6 mm dia., Kai Industries, Gifu, Japan), avoiding the puncture point of the syringe, and frozen in liquid nitrogen. qRT-PCR was performed according to the procedure described above using primers listed in Table S1. *NtActin* was used as the reference gene.

### *SiMYB2* genome structure

The selection of *SiMYB2* variants was investigated in several cultivars (AK, TK, GG, BB, MT, TR, SA, ST, KP) using RT-PCR. RNA extraction and RT-PCR were performed as described above. Reactions were performed at 98°C for 10 sec, 60°C for 5 sec, and 68°C for 1 sec for 40 cycles and amplified to a plateau. Primers were the same as those used for qRT-PCR (Table S1). *SiActin* was used as the internal control.

The genomic structure of *SiMYB2* was analyzed through Amplicon-seq using multiple lines and cultivars (AK, TK, GG, BB, MT, KP). First, genomic DNA was extracted from the leaves of these cultivars using the FavorPrep^TM^ Plant Genomic DNA Extraction Mini Kit (Favorgen). Primers were designed to amplify the ORF of *SiMYB2* (Table S1). PCR reactions were performed using KOD One^®^ PCR Master Mix (TOYOBO) under the following conditions: 98°C for 10 sec, and 68°C for 185 sec for 40 cycles. Subsequently, a library was prepared with the Native Barcoding Kit 24 V14 (SQK-NBD114.24, Oxford Nanopore Technologies, Didcot, UK) and sequenced on a FLO-MIN114 flow cell (Oxford Nanopore Technologies).

Based on the results of Amplicon-seq, primers were designed to detect insertions or deletions (In/Dels) within the *SiMYB2* locus (Table S1). Genomic DNA was extracted from the leaves of ST, SA, SH, and m-SH in addition to the cultivars listed above using a FavorPrep^TM^ Plant Genomic DNA Extraction Mini Kit (Favorgen). PCR reactions were performed using KOD One^®^ PCR Master Mix (TOYOBO). Reactions were performed at 98°C for 10 sec, 60°C for 10 sec, and 68 °C for 3 sec for 40 cycles.

The expression levels of *SiMYB2-Long* and *SiMYB2-Short* in the leaves of SH and m-SH were investigated by qRT-PCR. RNA extraction and qRT-PCR were performed as described above.

### Statistical analyses

All statistical processing was performed using R software (ver. 4.3.2, https://www.R-project.org/). Student’s *t*-test was performed using the ’stats’ package, and Dunnett’s range test was performed using the ’multcomp’ package.

## Results

### Variations in anthocyanin accumulation as a result of tissue culture

A total of 495 regenerated plants were obtained from KP leaf adventitious shoots, of which 96.6% (484/495) produced pink petals and reddish leaves, the same phenotype as KP (Table 1). 1.0% (5/495) did not accumulate anthocyanin and therefore had white petals and green leaves (KW). 0.8% (4/495) had randomly-colored petals, and 0.4% (2/495) had white-striped petals (KWS) (Fig. 1, Table 1). As the five KW plants were grown, two of them began to produce white striped petals like KWS, and the remaining three had randomly-pigmented petals (Fig. 1h, j). In addition, five KWs and two KWSs eventually grew petals and leaves like the parental KP (Fig. 1k). Fifty plants were regenerated from KW leaves, all of which had white petals and green leaves (50/50, Table 1). As these plants grew, some began producing striped or wholly-pigmented petals like those of KWS or KP, respectively.

**Table 1.**
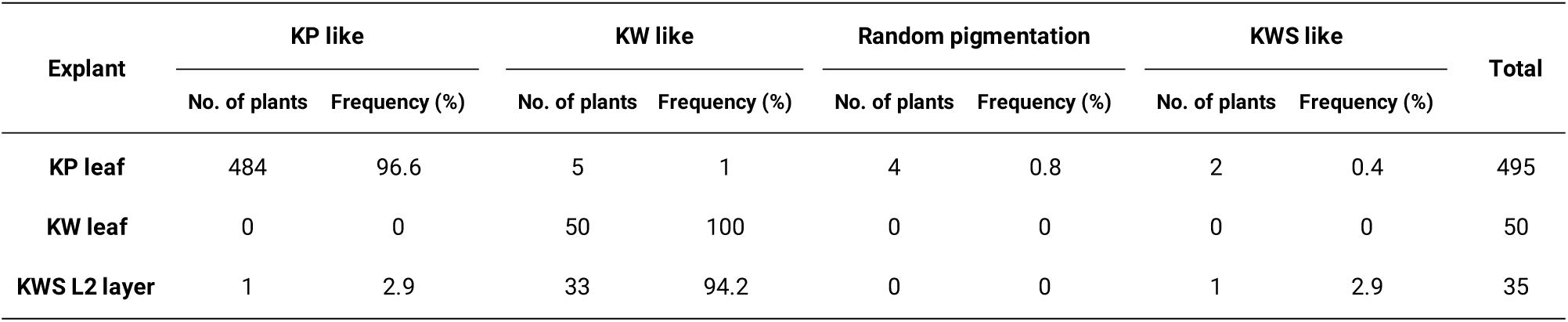
Investigation of the flower color phenotype of regenerated plants obtained by tissue culture.

If KWS is a periclinal chimera, plants derived from subepidermal tissue (L2 layer or below) should have only white petals and green leaves. On the other hand, if KWS is the result of epigenetic regulation, petal coloration may vary among individual plants. Based on this assumption, 35 plants were generated from subepidermal tissue. Petal coloring was distributed as follows: one plant (1/35) had pink petals, one (1/35) had white striped petals, and all the other plants had white petals (33/35, Table 1). As the plants grew, white petals began to produce pink pigment, white stripes, or randomly-colored petals. The other cultivar used in this study, ST, was also as unstable as ‘Kilauea’ for pigment patterning. For example, SW arising as a mutant from ST also became striped (SWS) and eventually had the same petal coloring as ST (Fig. 1i).

### Loss of anthocyanin biosynthesis in white petals and green leaves

Pelargonidin-3-acetylrutinose-5-glucoside (A1, Fig. S2a) and peonidin-3-acetylrutinose-5-glucoside (A2, Fig. S2a) were readily detected in KP petals, but were at low levels in KW petals (Fig. S2b). While apigenin-4’-glucuronide (F1, Fig. S2e) was identified as a flavone in KP petals, it was hardly detected in KW petals (Fig. S2f). In addition, cyanidin-3-acetylsambubioside (A3, Fig. S2c) and luteolin-4’-glucuronide (F2, Fig. S2g) were detected in KP leaves, but were not detected in KW leaves (Fig. S2d,h). Thus, the functions of *Chalcone synthase* (*CHS*), *Chalcone isomerase* (*CHI*), *Flavone synthase* (*FNS*), or all of them, may be disturbed in KW petals and leaves (Fig. 2a). Furthermore, total anthocyanin content was significantly lower in KW than KP petals and in SW than ST petals (Fig. 2a), and the anthocyanin content of KW leaves was lower than that of KP leaves (Fig. 2a).

**Fig. 2.**
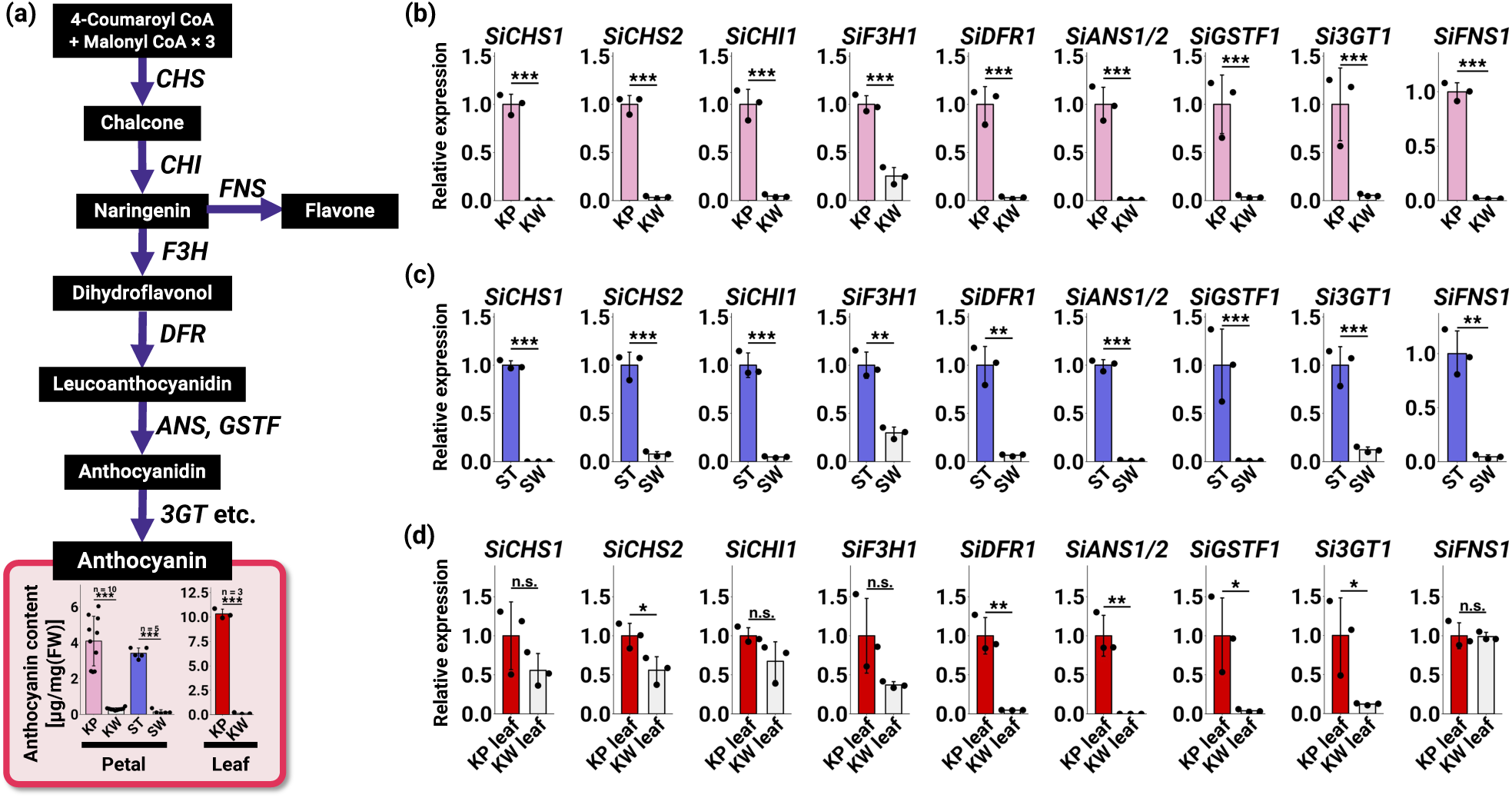
(a) Anthocyanin biosynthesis pathway and anthocyanin content in *Saintpaulia* petals and leaves. Asterisks indicate significant differences (Student’s *t*-test, \**P* < 0.05, \*\**P* < 0.01, \*\*\**P* < 0.001). n.s. indicates no significant difference. Error bars represent means ± SD (n = 10 for KP and KW petals, n = 5 for ST and SW petals, and n = 3 for KP and KW leaves). (b-d) qRT-PCR analysis of anthocyanin biosynthesis-related genes in petals. Asterisks indicate significant differences (Student’s *t*-test, \**P* < 0.05, \*\**P* < 0.01, \*\*\**P* < 0.001). n.s. indicates no significant difference. Error bars represent means ± SD (n = 3). (b) Expression in KP and KW petals. (c) Expression in ST and SW petals. (d) Expression in KP and KW leaves.

To investigate at which stage anthocyanin metabolism is interrupted, the expression levels of ABGs, *CHS* (*SiCHS1/2*), *CHI* (*SiCHI1*), and *FNS* (*SiFNS1*) in KP and KW petals were compared by qRT-PCR. The expression of *SiCHS1/2*, *SiCHI1,* and *SiFNS1* was significantly lower in KW petals than in KP petals (Fig. 2b). The expression levels of *SiF3H1*, *SiDFR1*, *SiANS1/2*, *SiGSTF1,* and *Si3GT1* were also significantly lower in KW petals than in KP petals (Fig. 2b). Consistent with the qRT-PCR results, RNA-seq showed that the expression levels of several ABGs (*SiCHS1/2*, *SiCHI1*, *SiF3H1*, *SiDFR*, *SiANS1/2*, *SiGSTF1*, *Si3GT1/2*, *SiMATE1/2/3*, etc.) were reduced in KW petals relative to KP petals (Table 2). SW petals, which have similar characteristics to KW, have several ABG expression levels significantly lower than in ST petals (Fig. 2c). In addition, the expression of several ABGs (*SiCHS1/2*, *SiCHI1*, *SiF3H1*, *SiDFR1*, and *SiANS1/2*) was lower in the white regions of KWS and SWS petals than in their pigmented regions (Fig. S3a, b). There were no significant differences in the expression levels of *SiCHS1*, *SiCHI1*, *SiF3H1,* and *SiFNS1* between KP and KW leaves (Fig. 2d), but the expression levels of *SiCHS2*, *SiDFR1*, *SiANS1/2*, *SiGSTF1*, and *Si3GT1* were significantly lower in KW leaves than in KP leaves (Fig. 2d). Based on qRT-PCR and RNA-seq results, the low anthocyanin accumulation levels in white petals and green leaves is likely due to the downregulation of several ABGs.

**Table 2.**
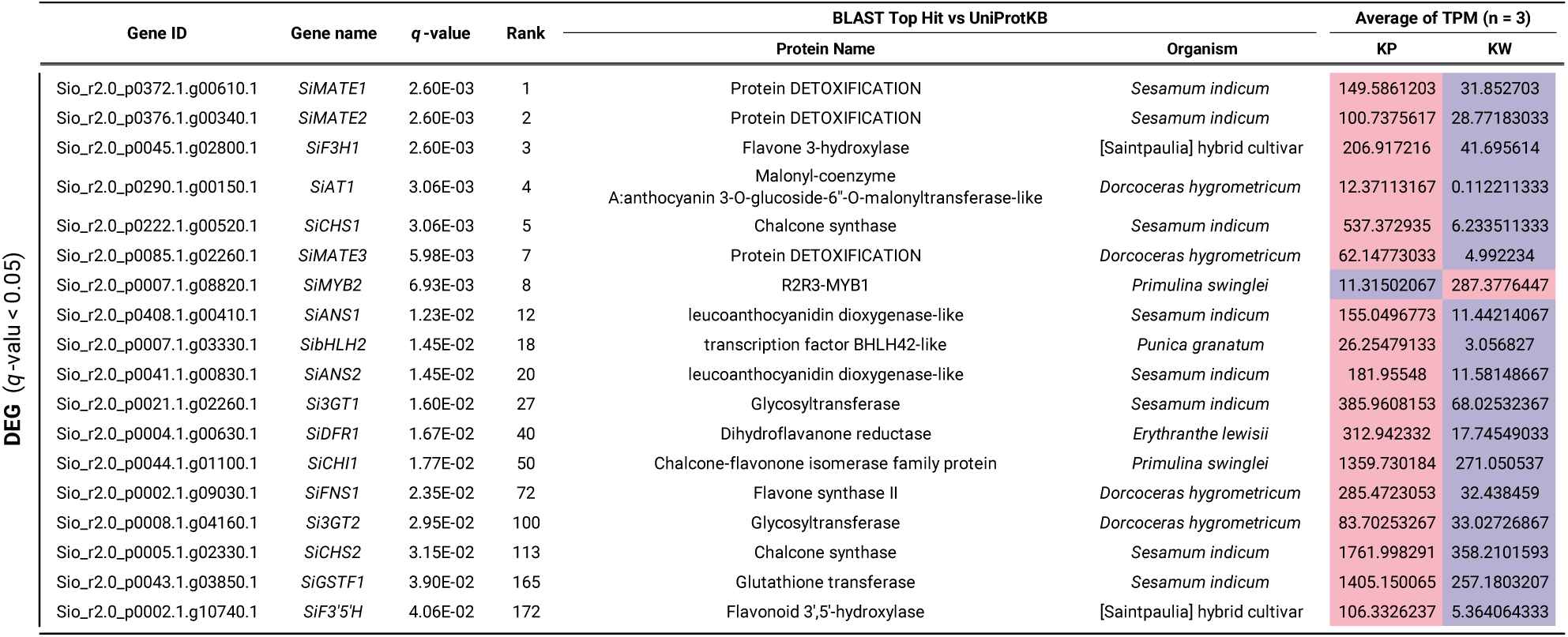
Summary of DEG analysis for anthocyanin biosynthesis related genes.

### Downregulation of anthocyanin biosynthesis-related transcription factors

The reduced expression of multiple ABGs in anthocyanin-deficient tissues suggests that TFs may be responsible for this downregulation, rather than transcript turnover or translation deficiencies. Based on the RNA-seq analysis of KP and KW petals, we focused on the key regulators of anthocyanin biosynthesis: *i.e. MYB*, *bHLH*, and *WDR* (Ramsay & Glover, 2005; Xu *et al*., 2015).

DEG analysis identified *SiMYB2* and *SibHLH2* as DEGs in petals, both of which are predicted to be TFs related to anthocyanin biosynthesis (Table 2). Expression levels of *SibHLH2* were significantly lower in KW than in KP petals, whereas *SiMYB2* expression levels were significantly higher in KW than in KP petals (Table 2). Consistent with the RNA-seq results, qRT-PCR indicated that *SibHLH2* expression was significantly lower in KW than in KP petals, and *SiMYB2* expression was significantly higher in KW than in KP petals (Fig. 3a). This expression pattern was also observed in a comparison between ST and SW petals (Fig. 3b). *SiMYB2* expression levels were also significantly higher in KW than in KP leaves, whereas *SibHLH2* expression levels were significantly lower in KW than in KP leaves (Fig. 3c), suggesting that either *SiMYB2*, *SibHLH2* or both are involved in the unstable pigmentation of *Saintpaulia* petals and leaves.

**Fig. 3.**
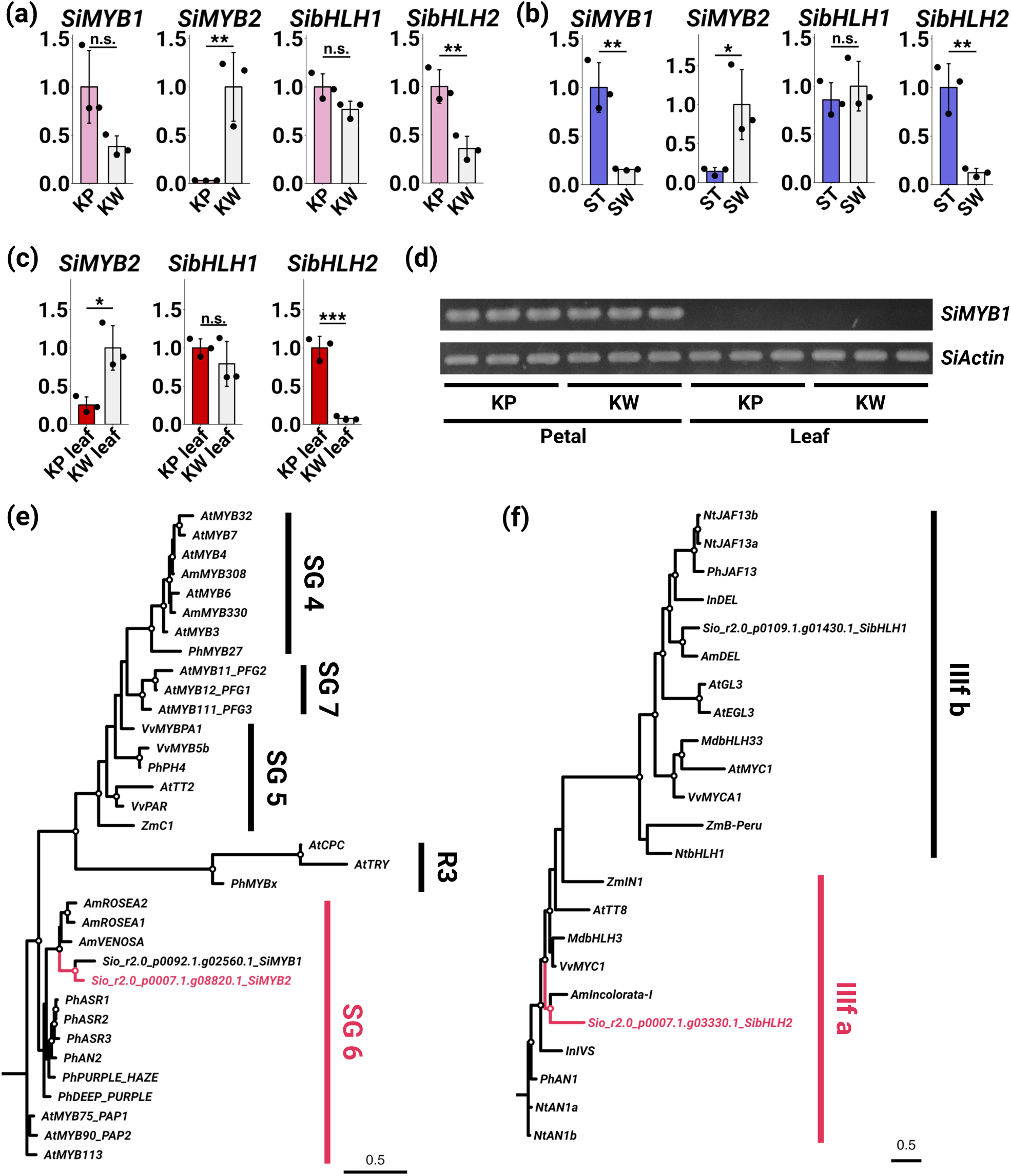
(a-c) qRT-PCR analysis of anthocyanin biosynthesis related transcription factors. Asterisks indicate significant differences (Student’s *t*-test, \**P* < 0.05, \*\**P* < 0.01, \*\*\**P* < 0.001). Error bars represent means ± SD (n = 3). (a) Expression levels in KP and KW petals. (b) Expression levels in ST and SW petals. (c) Expression levels in KP and KW leaves. (d) RT-PCR results of *SiMYB1* and *SiActin* in petals and leaves of KP and KW (n = 3). The amplification reaction was carried out for 35 cycles (plateau). (e-f) Molecular phylogenetic tree of MYB TFs (e) and bHLH TFs (f) amino acid sequences. Robust branches are indicated by white circles.

Although *SiMYB1* and *SibHLH1* were not DEGs, these TFs were anthocyanin biosynthesis-related ancillary to *SiMYB2* and *SibHLH2*. The qRT-PCR assay revealed that *SibHLH1* had equivalent expression levels in pigmented and non-pigmented tissues (Fig. 3a–c), and that *SiMYB1* expression was also about the same in KP and KW petals (Fig. 3a). In the non-pigmented mutant SW, which arose from ST, *SiMYB1* expression was significantly lower in SW than in ST (Fig. 3b). However, *SiMYB1* expression was nearly undetectable in the leaves of either KP or KW (Fig. 3d). These results suggest that *SiMYB1* and *SibHLH1* are unlikely to be the main factors of unstable anthocyanin accumulation.

A molecular phylogenetic tree was constructed to model the evolutionary history of MYB and bHLH TFs in *Saintpaulia*. The tree indicates that *SiMYB1* groups within the SG 6 clade (Stracke *et al*., 2001), which is known to promote anthocyanin biosynthesis (Fig. 3e). Although *SiMYB2* was highly expressed in non-pigmented tissues, and was hypothesized to belong to a clade associated with anthocyanin biosynthesis repression, it was instead clustered within the SG 6 clade (Fig. 3e). Both *SibHLH1* (IIIf b) and *SibHLH2* (IIIf a) belonged to clades known to promote anthocyanin biosynthesis (Heim, 2003) (Fig. 3f).

### Hypermethylation of *SiMYB2* in anthocyanin-deficient tissues

From the expression analysis results, *SiMYB2* and *SibHLH2* were identified as candidate genes involved in the unstable pigmentation of *Saintpaulia* petals and leaves. The hypothesis that loss and reversion of anthocyanin accumulation in flowers and leaves was associated with changes in the epigenome led us to consider that the epigenetic state of either *SiMYB2* or *SibHLH2* may have undergone modification. Therefore, WGBS (whole genome bisulfite sequence) was conducted to examine the methylation levels of these candidate genes in the leaves of KP and KW. Differences in methylation levels were observed in the second exon of *SiMYB2* (Fig. 4a). In particular, methylation levels in the CHG and CHH contexts were higher in KW than in KP (Fig. 4a, Table S3). Additionally, CHH context methylation was higher in KW in the promoter region of *SiMYB2* than in KP (Fig. 4a, Table S3). For *SibHLH2*, there were no differences in methylation levels in either the ORF or promoter regions (Fig. 4b, Table S3). The methylation levels of several other genes related to anthocyanin biosynthesis (*SiCHS1/2*, *SiCHI1*, *SiF3H1*, *SiDFR1*, *SiANS1/2*, *SiGSTF1*, *Si3GT1/2*, *Si3GRT1*, *SiMATE1/2/3*, *SiAT*, *SiF3’5’H*, *SiMYB1*, and *SibHLH1*) were also examined, but no differences were detected in the coding gene itself or its promoter regions (Fig. S4). McrBC-PCR/qPCR results also showed a difference in methylation levels in the second exon of *SiMYB2* between KP and KW leaves (Fig. 4c, f). This difference was also observed in KP and KW petals (Fig. 4d–f). Thus, *SiMYB2* was identified as an epiallele associated with unstable pigment accumulation.

**Fig. 4.**
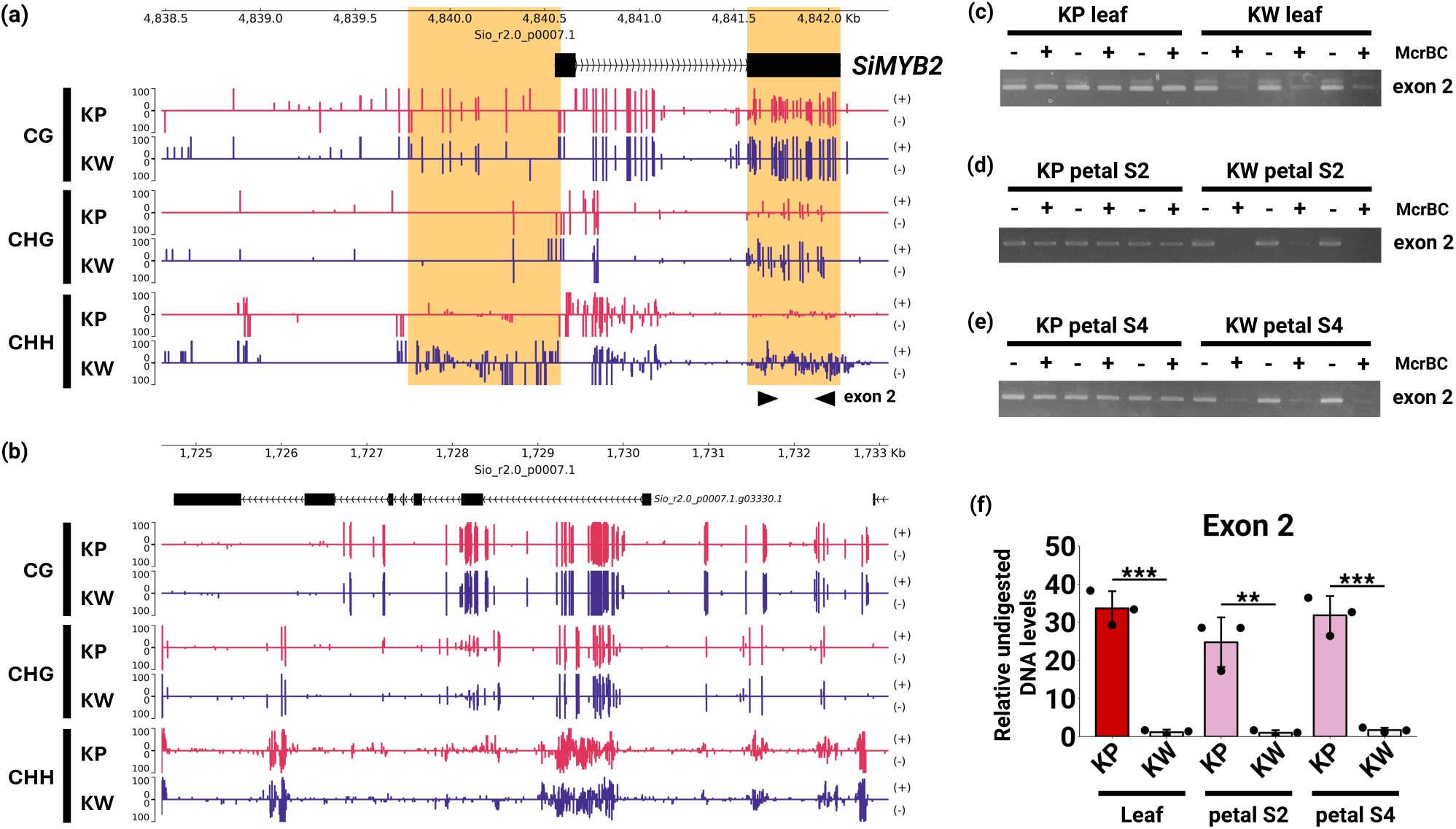
(a,b) Methylation levels of candidate genes responsible for unstable leaf pigmentation. The vertical axis indicates the methylation level, and the horizontal axis indicates the position in the Sio_r2.0_p0007.1 contig. Methylation levels are shown for each context (CG, CHG, CHH), and each strand (plus and minus). Red bars indicate methylation levels in KP, and blue bars in KW. (a) Methylation levels in *SiMYB2*. Regions where differences in methylation levels were observed are highlighted in orange. (b) Methylation levels in *SibHLH2*. (c-e) McrBC-PCR results of *SiMYB2* exon2 in leaf (c), petal at S2 (d), and petal at S4 (e) of KP and KW. (f) McrBC-qPCR results of *SiMYB2* exon2 in KP and KW tissues. Asterisks indicate significant differences (Student’s *t-test*, **P < 0.01, ***P < 0.001). The error bar represents mean ± SD (n = 3).

### Two transcripts selectively generated from the *SiMYB2* locus

*SiMYB2* (Sio_r2.0_p0007.1.g08820.1) was significantly up-regulated and accompanied by hypermethylation in anthocyanin-deficient tissues, suggesting its involvement in anthocyanin accumulation instability. Mapping of the *SiMYB2* genomic region showed a splice junction between *SiMYB2* and Sio_r2.0_p0007.1.g08810.1 in KP (Fig. 5a; sj2), suggesting that the *SiMYB2* gene region extends from Sio_r2.0_p0007.1.g08810.1 to Sio_r2.0_p0007.1.g08820.1. By Sanger sequencing of the TA cloning products and *de novo* assembly of RNA-seq reads, transcripts consisted of p1 + p2 + p4 and those of p3 + p4 were identified (Dataset S1). This result indicates that the *SiMYB2* locus is a single gene that ranges over approximately 18 kb, and produces at least two RNAs, presumably from two different transcription start sites (Fig. 5b, c). Exon 1 (124 bp), exon 2 (130 bp), exon 3 (110 bp), and exon 4 (493 bp) were determined from the sequences of p1, p2, p3, and p4 (Fig. 5b). The two 5’ UTRs (1 and 2) and one 3’ UTR were estimated. Transcripts consisting of 5’ UTR 1, exon 1, exon 2, exon 4, and 3’ UTR were designated as *SiMYB2-Long*, and 5’ UTR 2, exon 3, exon 4, and 3’ UTR as *SiMYB2-Short* (Fig. 5c). *SiMYB2-Long* was preferentially generated in KP and *SiMYB2-Short* in KW.

**Fig. 5.**
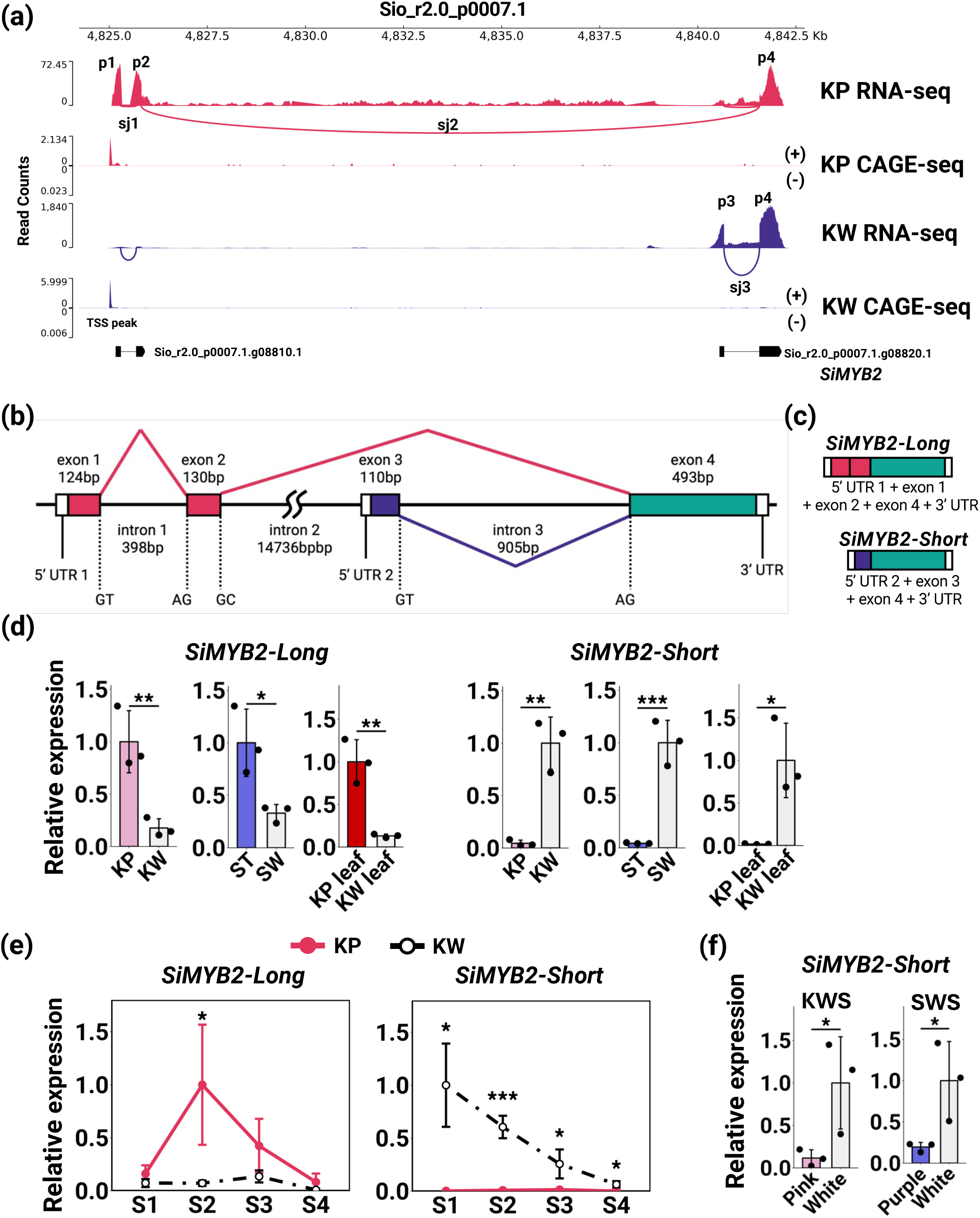
Characteristics of *SiMYB2*. (a) RNA-seq and CAGE-seq mapping within the *SiMYB2* locus (Sio_r2.0_p0007.1.g08820.1) and its upstream region. Vertical axes represent read counts. Horizontal axes represent a position on the contig Sio_r2.0_p0007.1. Peaks in RNA-seq indicate exon sequences. Splicing junctions (sj1/2/3) were located between p1, and p2, p2 and p4, and p3 and p4. Although p1 + p2 and p3 + p4 were predicted as independent genes (Sio_r2.0_p0007.1.g08810.1, Sio_r2.0_p0007.1.g08820.1), they were the same gene. Peaks in CAGE-seq indicate a transcription start site. (b-c) Expected structure of the *SiMYB2* locus based on mapping and cloning results. Transcripts consisting of exons 1, 2, and 4 were designated as *SiMYB2-Long*, and those consisting of exons 3 and 4 were designated as *SiMYB2-Short*. (d) qRT-PCR analysis of *SiMYB2* variants in KP and KW petals, ST, and SW petals, KP and KW leaves, pink and white regions in KWS petals, and purple and white regions in SWS petals. Asterisks indicate significant differences (Student’s *t*-test, \**P* < 0.05, \*\**P* < 0.01, \*\*\**P* < 0.001). Error bars represent means ± SD (n = 3).

The expression levels of *SiMYB2-Long* and *SiMYB2-Short* were confirmed by qRT-PCR. *SiMYB2-Long* expression levels were significantly lower in KW than in KP petals, and in SW than in ST petals (Fig. 5d). In contrast, *SiMYB2-Short* expression levels were significantly higher in KW than in KP petals, and in SW than in ST petals (Fig. 5d). This expression pattern for *SiMYB2-Long* and *SiMYB2-Short* was also observed in KP and KW leaves (Fig. 5d). The expression levels of *SiMYB2-Long* were significantly higher in KP compared to KW leaves, whereas the expression levels of *SiMYB2-Short* were significantly higher in KW than in KP leaves (Fig. 5d). The expression levels of *SiMYB2-Long* and *SiMYB2-Short* in KP and KW petals were also examined at different developmental stages. Anthocyanin accumulation was detected between S2–S3, with the expression levels of *SiMYB2-Long* being significantly higher at S2 in KP petals (Fig. 5e). In KW petals, the expression level was lower at all stages (Fig. 5e). *SiMYB2-Short* was hardly expressed at any stage in KP petals, while it was highly expressed in KW petals (Fig. 5e). Expression levels of *SiMYB2-Short* in the pigmented regions of striped flowers was significantly lower than in the non-pigmented regions of KWS and SWS petals (Fig. 5f), indicating that *SiMYB2-Long* is specifically expressed in colored tissues, whereas *SiMYB2-Short* is selectively transcribed in non-colored tissues.

*SiMYB2-Long* and *SiMYB2-Short* have different 5’ UTR sequences, suggesting that these variants arise from Alternative Promoter Usage (APU, Landry *et al*., 2003). CAGE-seq analysis was performed to determine the transcription start sites (TSSs) of *SiMYB2* variants. If the two variants were produced via APU, peaks would be expected to detect the upstream regions of exon 1 and exon 3 of *SiMYB2*, respectively. However, only a single TSS was detected upstream of exon 1 in both KP and KW samples (Fig. 5a). Consequently, the mechanism by which these two transcription variants are generated could not be determined in this study. Note that the upstream sequence of *SiMYB2-Short* contains a TA-rich region with several potential TATA boxes. (Fig. S5, Mukumoto et al., 1993).

### Functional analysis of *SiMYB2* by transient assays

The amino acid sequences of SiMYB1, SiMYB2-Long, and SiMYB2-Short were compared with other R2R3-MYB TFs belonging to the SG6 clade. SiMYB2-Short does not contain the entire R2 domain or most of the R3 domain, which are important conserved regions for MYB genes (Fig. 6a), but the R2-R3 domains of SiMYB1 and SiMYB2-Long were almost identical to those of other MYBs, and the bHLH binding domain ([D/E]Lx2[R/K]x3Lx6Lx3R) and SG 6 motif (KPRPR[S/T]F) were intact (Fig. 6a. Stracke et al., 2001; Zimmermann et al., 2004).

**Fig. 6.**
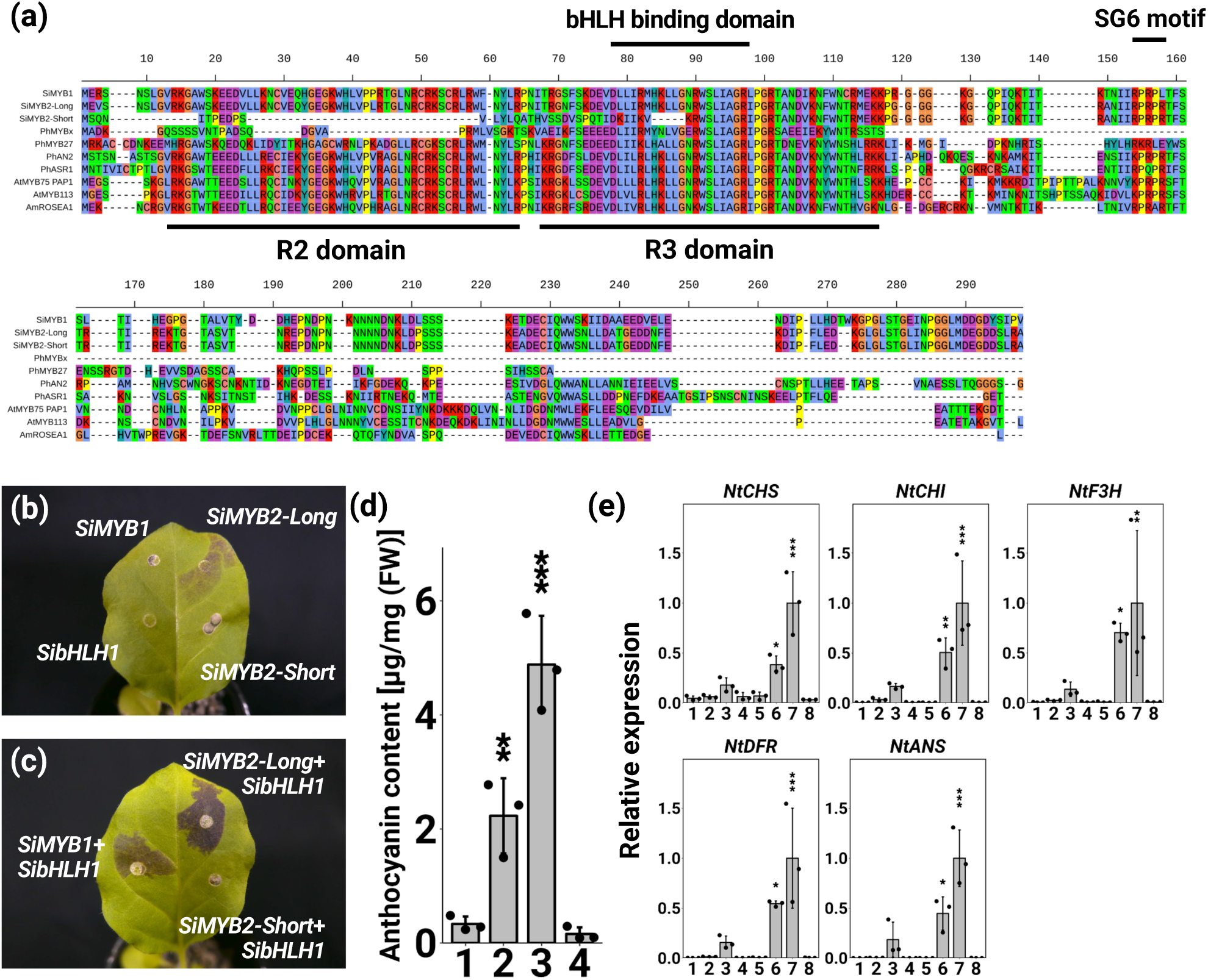
(a) Multiple alignment of MYB amino acids from *Saintpaulia* and other plants. The R2/R3 domain and SG 6 motif were based on Stracke et al. (2001), and the bHLH binding domain was based on Zimmermann et al (2004). (b-e) Results of transient assays using tobacco leaves. (b) Tobacco leaves when the target transcription factor (TF) was expressed alone. *SiMYB1* (top left), *SibHLH1* (bottom left), *SiMYB2-Long* (top right), and *SiMYB2-Short* (bottom right). (c) Tobacco leaves when MYB TFs were expressed simultaneously with *SibHLH1*. *SiMYB1* + *SibHLH1* (left), *SiMYB2-Long* + *SibHLH1* (top right), and *SiMYB2-Short* + *SibHLH1* (bottom right). Tobacco leaves were photographed four days post infiltration. (d) Anthocyanin content in tobacco leaves when MYB and bHLH TFs were expressed simultaneously. Anthocyanin content was determined as cyanidin-3-glucoside equivalents. Error bars represent means ± SD (n = 3). 1: GFP, 2: *SiMYB1* + *SibHLH1*, 3: *SiMYB2-Long* + *SibHLH1*, 4: *SiMYB2-Short* + *SibHLH1*. Asterisks indicate significant differences according to the Dunnett’s range test compared to *GFP*, (\**P* < 0.05, \*\**P* < 0.01, \*\*\**P* < 0.001). (e) qRT-PCR analysis of ABGs in infiltrated tobacco leaves. 1: GFP, 2: *SibHLH1*, 3: *SiMYB1*, 4: *SiMYB2-Long*, 5: *SiMYB2-Short*, 6: *SiMYB1* + *SibHLH1*, 7: *SiMYB2-Long* + *SibHLH1*, 8: *SiMYB2-Short* + *SibHLH1*. Error bars represent means ± SD (n = 3). Asterisks indicate significant differences according to the Dunnett’s range test compared to *GFP*, (\**P* < 0.05, \*\**P* < 0.01, \*\*\**P* < 0.001).

The functions of SibHLH1, SiMYB1, SiMYB2-Long and SiMYB2-Short were examined by transient assays in tobacco leaves. Active expression of the gene of interest was confirmed by qRT-PCR (Fig. S6). Expression of *SiMYB2-Long* alone resulted in very little accumulation of anthocyanins (Fig. 6b). When *SibHLH1*, *SiMYB1*, or S*iMYB2-Short* were expressed individually, no anthocyanin accumulation was observed (Fig. 6b). Subsequently, *SibHLH1* was selected as a potential cofactor for *SiMYB1* and *SiMYB2*, as its expression levels did not differ between colored and non-colored tissues. Anthocyanin accumulated when *SiMYB1* plus *SibHLH1*, or *SiMYB2-Long* plus *SibHLH1* were expressed (Fig. 6c). There was no accumulation of anthocyanin in tissues where *SiMYB2-Short* was co-expressed with *SibHLH1* (Fig. 6c). Anthocyanin content was also measured when MYB and bHLH were simultaneously expressed. The anthocyanin content of *SiMYB2-Long* co-expressed with *SibHLH1* was the highest among all transient assays, while anthocyanin content of tissues co-expressing *SiMYB2-Short* and *SibHLH1* was not different from the GFP control (Fig. 6d). The anthocyanin content of *SiMYB1* plus *SibHLH1* was lower than that of *SiMYB2-Long* plus *SibHLH1*, as reflected in its fainter coloring (Fig. 6d). The expression levels of ABGs were confirmed when these TFs were expressed. Several ABGs were upregulated in *SiMYB2-Long* only, *SiMYB1* plus *SibHLH1*, and *SiMYB2-Long* plus *SibHLH1*, whereas there was no upregulation of ABGs in S*iMYB2-Short* plus *SibHLH1* co-expression assays (Fig. 6e). Compared to *SiMYB1* plus *SibHLH1*, *SiMYB2-Long* plus *SibHLH1* had higher ABG expression, which is consistent with its anthocyanin content (Fig. 6e).

To investigate the possibility that SiMYB2-Short, whose expression is higher in non-pigmented tissues, indirectly inhibits R2R3-MYB function in a dosage-dependent manor, similar to CPC type R3-MYB (Li *et al*., 2020), bacterial suspensions expressing gradually increasing ratios of SiMYB2-Short to SiMYB1 and SibHLH1 were injected into tobacco leaves. Anthocyanin accumulation was not affected by changing the ratio of SiMYB2-Short (Fig. S7).

### The origin of *SiMYB2-Short* is the insertion of a transposon-like sequence

RT-PCR was performed to investigate whether the transcriptional selection of *SiMYB2-Long* and *SiMYB2-Short* also occurs in other cultivars. Two *SiMYB2* variants were found to be transcribed in TR and SA, which are similar to KP and ST (Fig. S8j). In contrast, *SiMYB2-Long* was amplified in AK, TK, GG, BB, and MT, but *SiMYB2-Short* was not (Fig. S8j). TR and SA, like KP and ST, are cultivars in which anthocyanin accumulation is unstable, showing loss and reversion during tissue culture propagation (Fig. S8f-i). AK, TK, GG, BB, and MT are cultivars with consistent anthocyanin accumulation in their leaves and flowers (Fig. S8a-e).

The genomic structure of *SiMYB2* was examined in AK, TK, GG, BB and MT using Amplicon-seq, which showed that there is a ∼2.5 kb deletion, including exon 3, which is required for the expression of *SiMYB2-Short*, in all five cultivars (Fig. 7a, S7k). To validate this finding, genomic PCR was performed using primers designed to flank the deletion region (Fig. 7a). As with the Amplicon-seq results, only the deletion-type band was amplified in AK, TK, GG, BB, and MT (Fig. 7b). In KP, only the insertion-type band was amplified, whereas in ST, TR, and SA, both insertion- and deletion-type bands were detected (Fig. 7b). These results indicate that the absence of *SiMYB2-Short* expression in AK, TK, GG, BB, and MT is due to deletion.

**Fig. 7.**
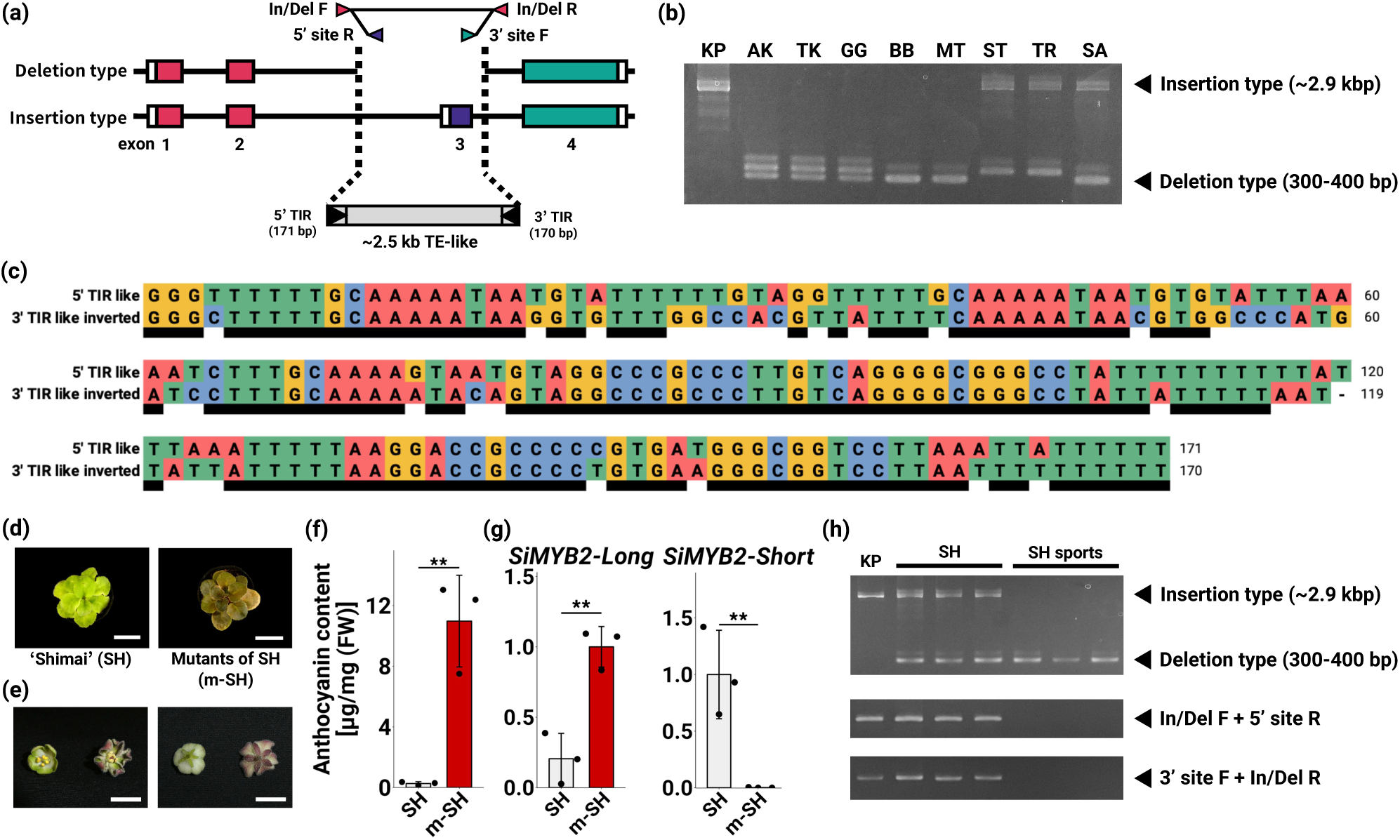
(a) The position of primers designed from the *SiMYB2* locus sequence is indicated by arrowheads. (b) Confirmation of indels by PCR in some cultivars. The abbreviated names of the cultivars are shown across the top of the gel lanes. The positions of the primers (In/Del F and R) used are shown in (a). (c) Multiple sequence alignment results of TIR-like sequences. Black bars indicate corresponding bases between 5’ TIR and 3’ TIR inverted sequences. (d-h) Characterization of the mutant obtained by tissue culture (m-SH) and parental cultivar (SH). (d) Photo of rosettes. White bars indicate 5 cm scale. (e) Photo of flowers. White bars indicate 1.5 cm scale. (f) Anthocyanin content of SH and m-SH leaves. The content was determined as cyanidin-3-glucoside equivalents. Error bars represent means ± SD (n = 3). Asterisks indicate significant differences (Student’s *t*-test, \**P* < 0.05, \*\**P* < 0.01, \*\*\**P* < 0.001). (g) qRT-PCR quantification of SiMYB2 variants in leaves. Error bars represent means ± SD (n = 3). Asterisks indicate significant differences (Student’s *t*-test, \**P* < 0.05, \*\**P* < 0.01, \*\*\**P* < 0.001). (h) Confirmation of the presence of a transposon-like element by PCR in SH and m-SH. The positions of the primers used are shown in (a).

The deleted region DNA sequence contains features of a transposon-like element (TE-like), including terminal inverted repeats (TIRs) (Fig. 7a,c). The 5’ TIR was 171 bp, the 3’ TIR was 170 bp, and the target site duplication (TSD) was 10 bp. A BLAST search for homologous transposon sequences failed to reveal any highly similar matches. Using the TIR sequences as queries, a BLAST search against the *Saintpaulia* SIO_r2.0 genome identified numerous homologous sequences (Table S4), suggesting that *SiMYB2-Short* may have arisen due to the insertion of an endogenous TE-like element.

Like KW and SW, SH is a cultivar that does not accumulate anthocyanins in its leaves. When SH was propagated via tissue culture, the resulting plants also did not accumulate anthocyanin in their leaves, like the original SH phenotype (Fig. 7d), and flowers of regenerated plants are green and do not accumulate anthocyanins (Fig. 7e). However, a single mutant, referred to as m-SH, emerged with red-tinted leaves and flowers (1 of 50 regenerated plants. Fig. 7d,e). The anthocyanin content of m-SH leaves was significantly higher than in wild type SH leaves (Fig. 7f). The expression of *SiMYB2-Long* was barely detectable in SH, but was highly expressed in m-SH (Fig. 7g). *SiMYB2-Short* was expressed in SH but was not detected in m-SH (Fig. 7g).

The genomic structure of *SiMYB2* was examined in these lines using In/Del primers, as in previous analyses. SH showed amplification of both insertion- and deletion-type bands (Fig. 7h), whereas in m-SH, only the deletion-type band was detected (Fig. 7h). These results suggest that the presence of *SiMYB2-Short* expression or the TE-like region may be essential for the loss of anthocyanin accumulation or its instability.

## Discussion

In this study, we identified an epiallele involved in unstable pigmentation patterning in *Saintpaulia* plants. Anthocyanins did not accumulate, and the expression levels of several ABGs were low in non-pigmented tissues. RNA-seq analysis was performed to determine candidate TFs, and a MYB TF, *SiMYB2*, was identified as an epiallele through WGBS analysis. Two transcripts were generated from the *SiMYB2* locus, one of which was highly expressed in pigmented colored tissues, and the other in non-pigmented tissues. These transcripts were designated as *SiMYB2-Long* and *SiMYB2-Short*, respectively, reflecting their relative lengths. Transient assays showed that *SiMYB2-Long* is an anthocyanin biosynthesis activator. *SiMYB2-Short* was highly expressed in non-pigmented tissues, suggesting that it is a repressor-type MYB TF. Paradoxically, the transient assay results also revealed that *SiMYB2-Short* does not have either a positive or negative impact on anthocyanin expression. Sequence analysis of the *SiMYB2* gene among several cultivars suggested that *SiMYB2-Short* resulted from the insertion of a transposon-like sequence. These observations provide a compelling model to help explain why mutants generated by tissue culture from an anthocyanin-deficient cultivar in which *SiMYB2-Short* is expressed accumulated anthocyanins.

### Formation of white striped petals due to periclinal chimeras versus epigenetics

The white striped flowers produced by *Saintpaulia* are thought to be the result of a periclinal chimeric structure (Lineberger & Druckenbrod, 1985; Kazemian *et al*., 2019). However, there is currently no evidence that they are due to the peripheral chimera structure. In this study, the tissue culture progeny of *Saintpaulia* KP and KWS flowers varied in their petal and leaf color patterns (Fig. 1a, Table 1). A previous study reported that the phenotypic variation of petal color varied greatly in plants regenerated from the tissue culturing of cultivar ‘Kilauea’ (Yang *et al*., 2017). The characteristic of *Saintpaulia* phenotype that changes in tissue culture and then reverts to the original phenotype suggests that this phenotype is the result of changes in epigenetic regulation.

White-striped petals closely resemble striped petals caused by periclinal chimeras, which may lead to the misconception that they too are periclinal chimeras. Periclinal chimeras occur when the cell layers of the shoot apical meristem (SAM) have distinct genetic backgrounds and coexist within a single plant (Satina, 1940). When such genetic differences involve anthocyanin biosynthesis-related genes, *Saintpaulia* flowers can exhibit striped petals. In this study, *Saintpaulia* with white striped petals eventually flowered with fully colored petals. Moreover, white-striped petals arose post-developmentally from *Saintpaulia* plants which had initially exhibited entirely white petals. Thus, it is unlikely that white-striped petals are due to periclinal chimeras. How, then, are the white-striped petals formed? In white-striped petals, the presence or absence of *SiMYB2-Short* expression correlated to lack of pigmentation (Fig. 5g). Additionally, the transcriptional selection of *SiMYB2* variants appears to be regulated epigenetically, suggesting that white-striped petals may result from cell layers with distinct epigenetic backgrounds. In other words, it may be an epigenetic periclinal chimera. Indeed, genes with cell layer-specific expression have been reported in *A. thaliana* (Lu *et al*., 1996). The epigenomic states of the L2 layer or below may be altered by tissue culture (Fig. 8). In white-striped petals, the epigenomic states of different cell layers in SAM may be independently regulated, allowing these distinct states to coexist within the same tissue (Fig. 8).

**Fig. 8.**
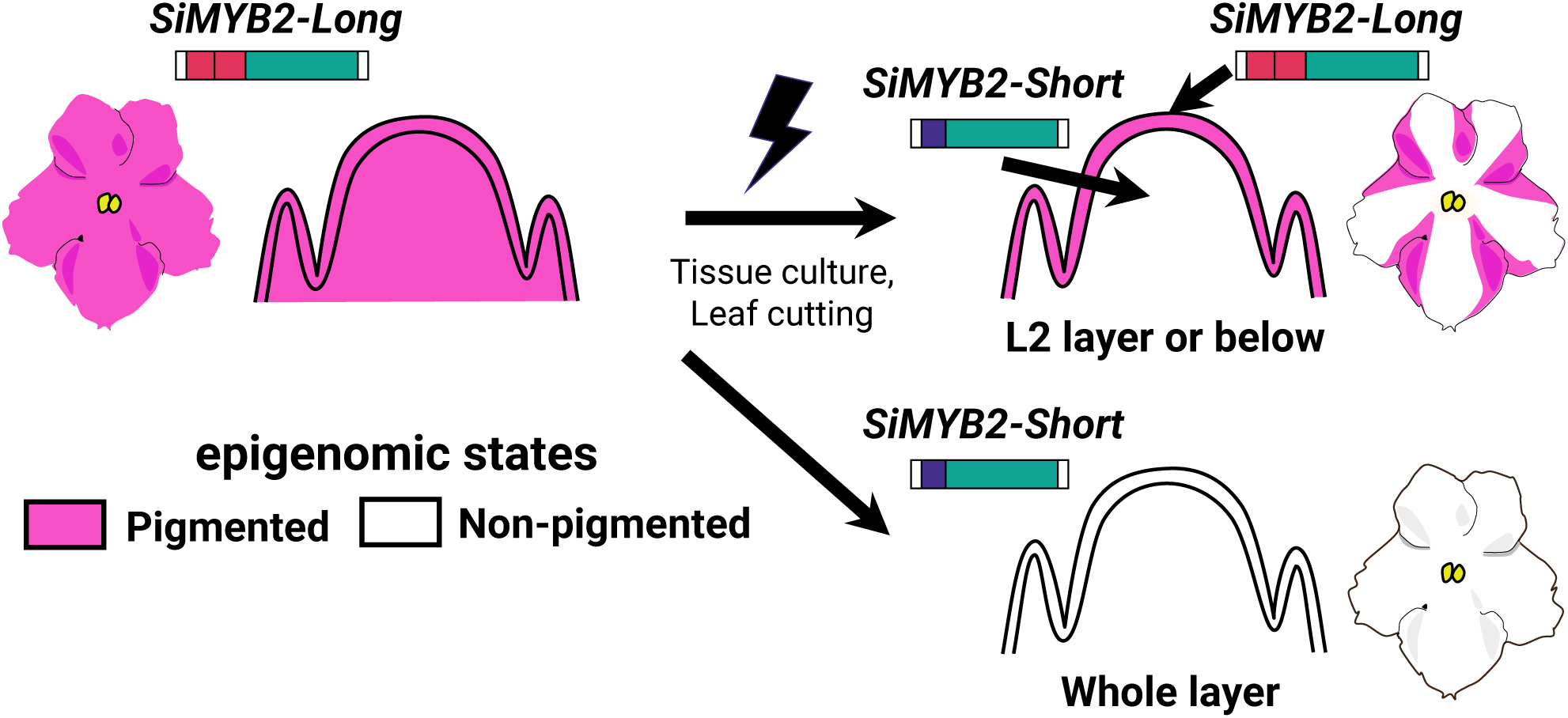
Hypothetical model to explain white stripe patterns. Pigmented petals have an epigenomic state in which *SiMYB2-Long* is expressed. Non-pigmented petals have an epigenomic state in which *SiMYB2-Short* is expressed. White striped petals may be the result of an epigenomic state in which *SiMYB2-Long* is expressed in the L1 layer and *SiMYB2-Short* in the L2 layer and below.

### Identification of the epiallele linked to unstable pigmentation

The traits of plants obtained through tissue culture or vegetative propagation are not always consistent. Such variations are occasionally attributable to alterations in epigenetic status. For example, during the tissue culture of *E. guineensis*, a reduction in the methylation level of the *Karma* transposable element within the *Deficiens1* (*DEF1*) gene leads to splicing errors in the *DEF1* gene, resulting in malformed fruit (Ong-Abdullah *et al*., 2015). In *C. morifolium*, yellow-flowered plants arise from the lateral buds of red-flowered plants. This phenomenon is caused by hypermethylation in the promoter region of *CmMYB6* (Tang *et al*., 2022). In our case, tissue culture may have altered the epigenetic state of *Saintpaulia*, leading to phenotypic variations.

Based on the hypothesis that unstable anthocyanin accumulation in *Saintpaulia* is caused by epigenetic regulation, we determined that *SiMYB2* is the epiallele associated with coloration by a combination of RNA-seq and WGBS analyses. *CmMYB6* in *C. morifolium* was identified as an epiallele involved in flower color changes using a similar approach as in the present study (Tang *et al*., 2022). *EgDEF1* in *E. guineensis* was also identified by epigenome-wide association analysis (EWAS) and WGBS (Ong-Abdullah *et al*., 2015). Exploration of other epialleles that control unstable traits, such as unstable anthocyanin accumulation, by this combination of methods may yield additional useful candidates for the identification of causal genes. *Saintpaulia* has been propagated and maintained in large quantities by vegetative propagation such as leaf cuttings and tissue culture. Epigenetic mutants, such as the anthocyanin-depleted individuals in this study, are often produced by vegetative propagation that are maintained until the next generation. For example, variation in transposon transposition frequency and altered floral patterns have been reported in tissue culture progeny of *Saintpaulia* ’Thamires’ (Sato *et al*., 2011). It is also postulated that these traits are the result of epi-mutation. It may therefore be feasible to utilize the epigenetic traits observed in *Saintpaulia* for genetic identification purposes.

The present study identified *SiMYB1* and *SibHLH2* as TFs associated with anthocyanin accumulation in petals, in addition to *SiMYB2* (Fig. 3). The transient assay results demonstrated that *SiMYB1* functions as an anthocyanin biosynthesis activator (Fig. 6c-e). Therefore, *SiMYB1* expression may be the determinant of unstable pigmentation. However, *SiMYB1* was barely expressed in leaves exhibiting unstable pigment accumulation like petals (Fig. 3d). In contrast, the transcriptional selectivity of *SiMYB2-Long* and *SiMYB2-Shor*t was linked to pigment accumulation in the leaves (Fig. 3c, 7f). These results indicate that *SiMYB1* is not the gene responsible for the phenomenon of unstable pigmentation.

The expression levels of *SibHLH1*, which is thought to regulate the transcription of ABGs, were essentially the same in white and colored petals. Phylogeny placed *SibHLH1* in the IIIf b clade, which includes *PhJAF13* (Fig. 3f. Quattrocchio et al., 1998; Heim, 2003). In Petunia, the expression of *PhJAF13* is not influenced by the expression of MYB TFs (Quattrocchio *et al*., 1998; Spelt *et al*., 2000), which is consistent with our observation that the expression of *SibHLH1* was unchanged in the white parts of *Saintpaulia*, where the expression of the MYB TFs was downregulated. *SibHLH2* clustered with the IIIf a clade, which includes *PhAN1* (Fig. 3f. Spelt et al., 2000; Heim, 2003). The loss of *PhAN1* function results in the loss of anthocyanin biosynthesis in Petunia (Spelt *et al*., 2000). Thus, the reduction of *SibHLH2* expression may contribute to the loss of anthocyanin in *Saintpaulia*. It has been reported that *NtJAF13*, a member of the IIIf b clade, interacts with MYB TFs and promotes the expression of *NtAN1*, a member of the IIIf a clade in *N. tabacum* (Montefiori *et al*., 2015). Moreover, *AmDelila*, a member of clade IIIf b, binds to the *Rosea1* MYB TF and positively regulates the transcription of *AmIncolorata I*, a member of clade IIIf a, in *Antirrhinum majus* (Albert *et al*., 2021). Therefore, the downregulation of *SibHLH2* in the non-pigmented tissues is most likely due to decreased *SiMYB2-Long* expression levels. Moreover, *SiMYB2* methylation levels differed between KP and KW, while *SibHLH2* did not (Fig. 4), suggesting that differences in *SibHLH2* expression are not the underlying cause of unstable pigmentation.

### Transcriptional selectivity of *SiMYB2* variants and pattern formation

Alternative splicing (AS) and APU are two regulatory mechanisms by which different transcripts can be produced from a single locus (Reviewed in Landry et al., 2003). An example of unstable pigmentation caused by different variants has been reported in *C. morifolium*. AS of *CmbHLH2* produces different variants, with non-functional *CmbHLH2* in white petals and functional *CmbHLH2* in colored petals (Lim *et al*., 2021). However, the present work is the first report that two or more transcripts from a single locus are involved in flower pattern formation. It was thought that *SiMYB2* variants were generated by APU since the 5’ UTRs of *SiMYB2-Long* and *SiMYB2-Short* were unique to each, particularly since multiple TATA boxes were observed in the predicted promoter region of *SiMYB2-Short*. However, we were unable to detect the TSS of *SiMYB2-Short* by CAGE-seq, thus preventing a conclusive determination about whether *SiMYB2* variants are generated through APU. CAGE-seq specifically sequences the 5’ ends of RNAs with a 5’ cap structure (m7G). Therefore, the absence of a detectable TSS for *SiMYB2-Short* may be attributed to the lack of a 5’ cap structure in *SiMYB2-Short* transcripts. Although this study could not identify the precise mechanism by which *SiMYB2* variants are generated, the findings from CAGE-seq provide valuable insights for future *SiMYB2* variant studies.

*SiMYB2-Short* was preferentially transcribed in both petals and leaves where anthocyanins were absent, and was already highly expressed by stage S1 in KW petals, which was earlier than the developmental stage at which *SiMYB2-Long* was expressed in KP petals. Mutants lacking the TE-like sequence accumulated anthocyanins at high levels. In other anthocyanin-accumulating cultivars, the loss of the TE-like sequence may result in anthocyanin production because transcription of functional *SiMYB2-Long* is the only possibility. However, anthocyanin accumulation can still occur in some cases even in the presence of the TE-like sequence. Notably, in tissues where anthocyanins accumulate, *SiMYB2-Short* is not transcribed, while *SiMYB2-Long* was actively transcribed. These findings suggest that non-production of anthocyanin may result from the transcription of *SiMYB2-Short* rather than the mere presence of the TE-like sequence.

The specific function of *SiMYB2-Short* could not be identified in this study, so further research is necessary to determine its role. If stable anthocyanin accumulation is desired, deleting the TE-like sequence may be useful. On the other hand, the presence of the TE-like sequence may enable transcriptional selectivity between *SiMYB2* variants, potentially giving rise to patterned flowers, albeit with instability. It is important to note that the presence of the TE-like insertion alone does not necessarily result in anthocyanin loss; additional factors may influence the transcriptional selectivity of *SiMYB2-Long* and *SiMYB2-Short*.

## Acknowledgments

We are grateful to Dr. Atsushi Hoshino of the National Institute for Basic Biology for his guidance in identifying transposons. We also wish to acknowledge the assistance provided by Assistant Professor Akira Yamazaki of Kindai University in the preparation and use of experimental equipment. We also thank Mr. Ryutaro Kunimune and Kazuki Kobayashi of Kindai University for their help in cultivation management. We would like to express our gratitude to Dr. Yu Kinoshita of Kyoto University for his invaluable guidance on the subject of epigenetics and for her assistance with cultivation. This work partly was supported by Sasakawa Scientific Research Grant from the Japan Science Society. Computations were partially performed on the NIG supercomputer at ROIS National Institute of Genetics.

## Competing Interest Statement

The authors have declared no competing interest.

## Author contributions

DK: Data curation, Investigation, Resources, Software, Visualization, Formal analysis, Conceptualization, Project administration, Writing-original draft. TT: Data curation, Investigation, Resources KS: Funding acquisition, Data curation, Formal analysis, Writing-review & editing. HH: Funding acquisition, Data curation, Formal analysis, Writhing-review & editing. FT: Data curation, Investigation, Writhing-review & editing. MH: Funding acquisition, Resources, Supervision, Project administration, Conceptualization, Writing-review & editing.

---

**Fig. S1.**
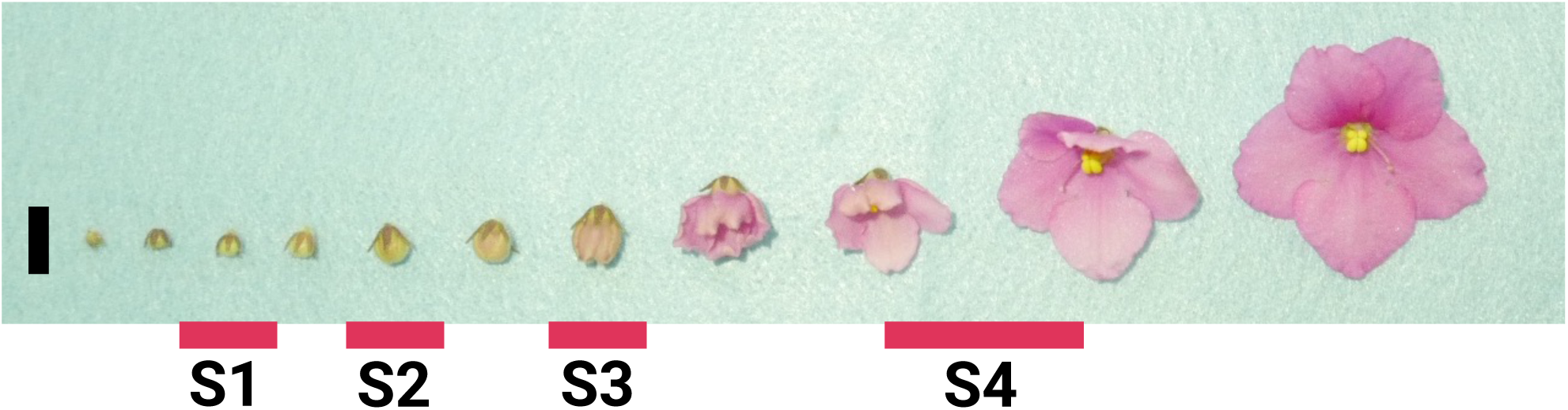
Developmental stages of petals used in this study. Black bar indicates 1 cm scale.

**Fig. S2.**
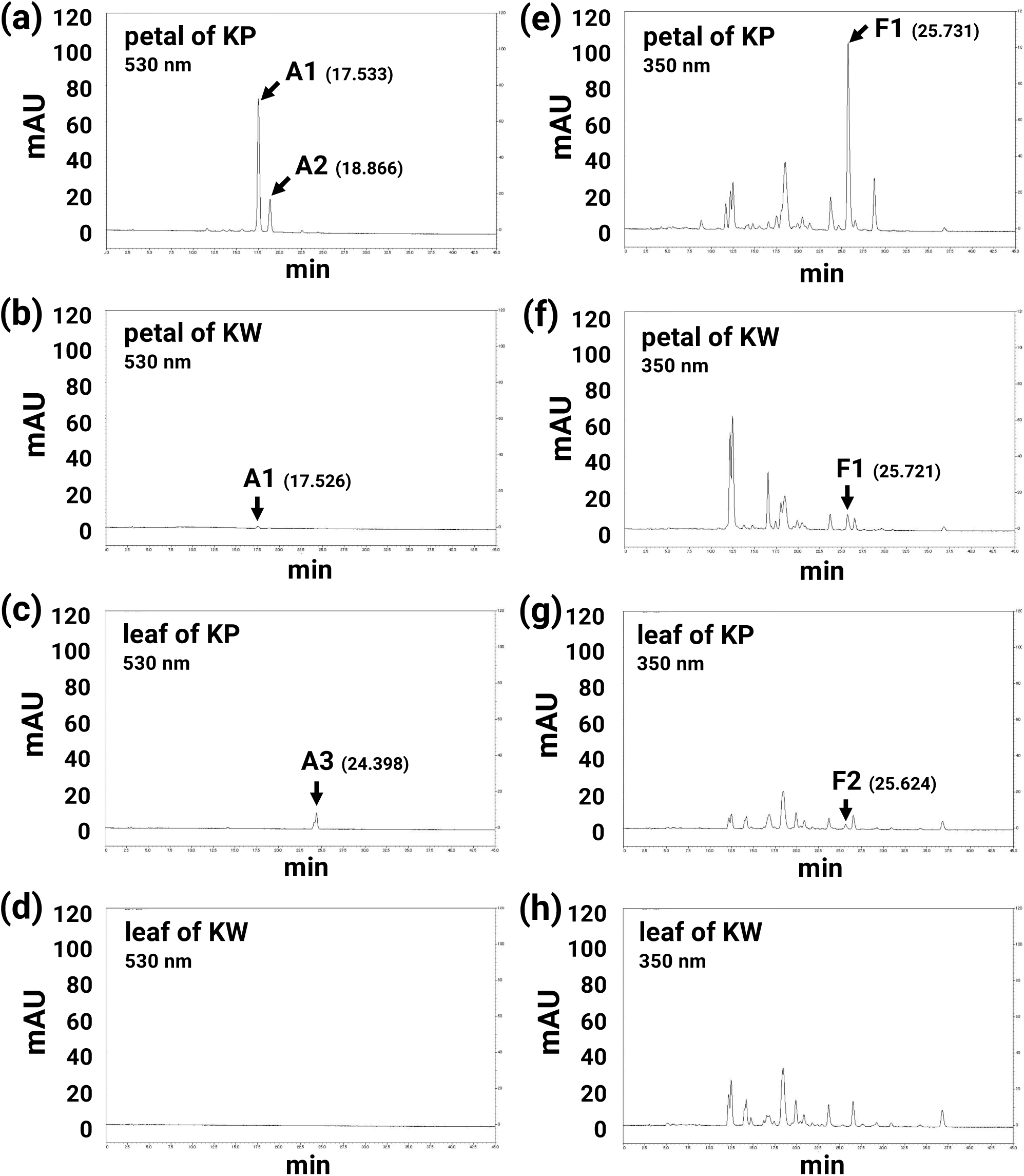
HPLC analysis of KP and KW petals. (a,b,e,f) Petal. (c,d,g,h) Leaf. (a–d) 530 nm for anthocyanin. (e–h) 350 nm for flavone. A1: Pelargonidin-3-acetyl-rutinoside-5-glucoside, A2: Peonidin-3-acetyl-rutinoside-5-glucoside, A3: Cyanidin-3-acetyl-sambubioside, F1: Apigenin-4’-glucronide, F2: Luteolin-4’-glucronide.

**Fig. S3.**
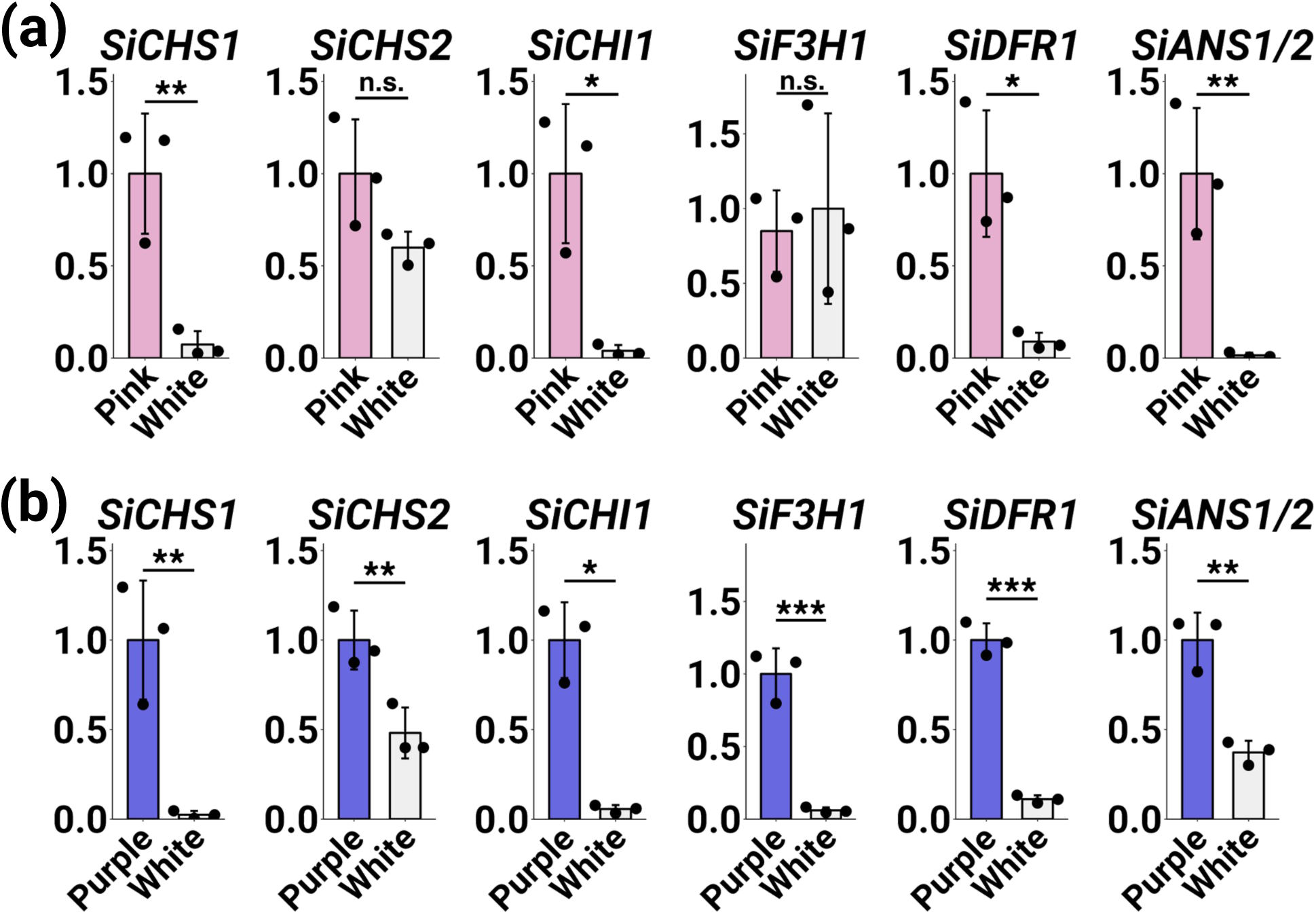
Expression levels of ABGs in white-striped petals. Asterisks indicate significant differences according to Student’s *t*-test (\**P* < 0.05, \*\**P* < 0.01, \*\*\**P* < 0.001). n.s. indicates no significant difference. Error bars represent means ± SD (n = 3). (a) Expression levels in KWS petals. Pink indicates pink regions in KWS petals. White indicates white regions. (b) Expression levels in SWS petals. Purple indicates purple regions in SWS petals. White indicates white regions.

**Fig. S4.**
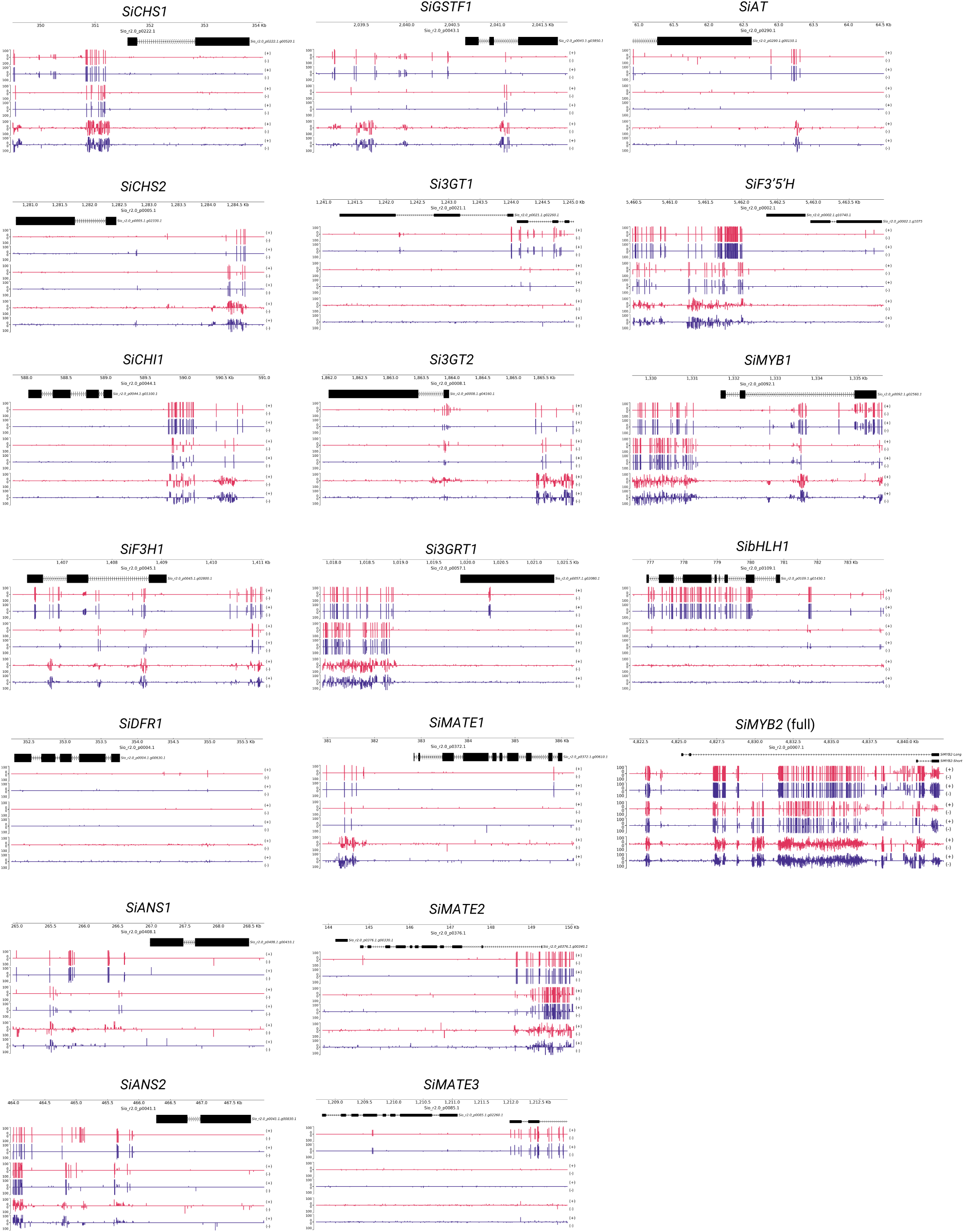
Methylation levels of anthocyanin biosynthesis related genes in KP and KW leaves.

**Fig. S5.**
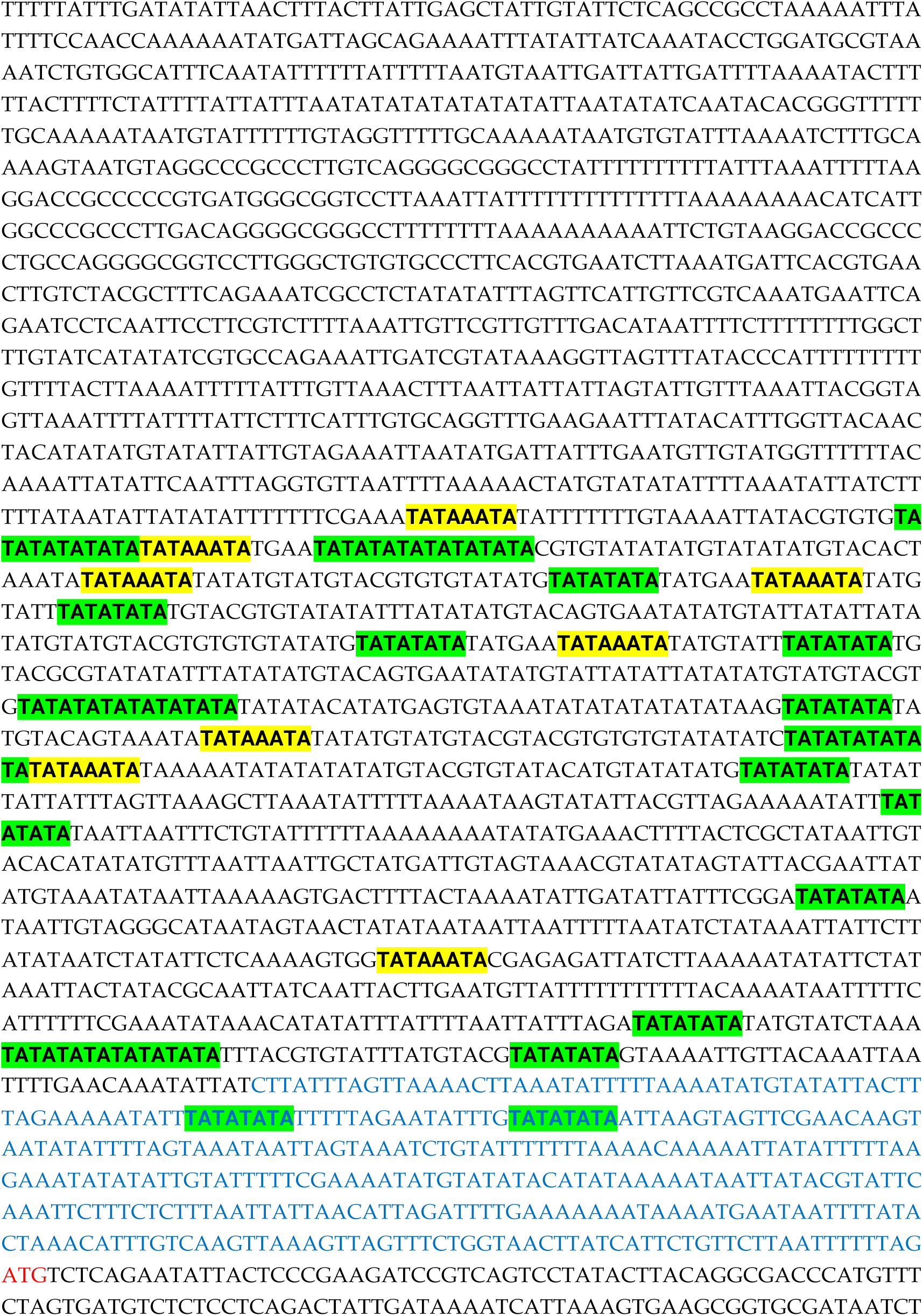
Predicted promoter sequences of *SiMYB2-Short*. Two types of TATA-boxes reported by Mukumoto et al. (1993) are highlighted. Start codon is indicated in red letters. Predicted 5’ UTR sequences is indicated in blue letters.

**Fig. S6.**
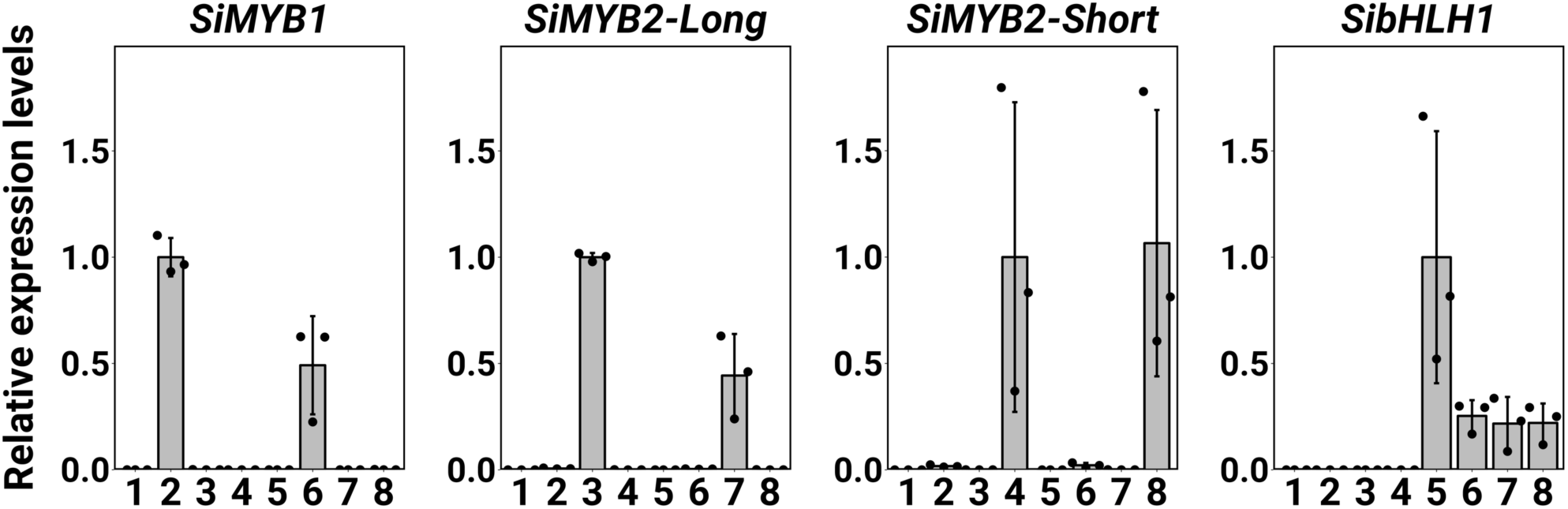
qRT-PCR analysis of the agroinfiltrated genes in tobacco leaves. 1: GFP, 2: *SibHLH1*, 3: *SiMYB1*, 4: *SiMYB2-Long*, 5: *SiMYB2-Short*, 6: *SiMYB1* + *SibHLH1*, 7: *SiMYB2-Long* + *SibHLH1*, 8: *SiMYB2-Short* + *SibHLH1*. Error bars represent means ± SD (n = 3).

**Fig. S7.**
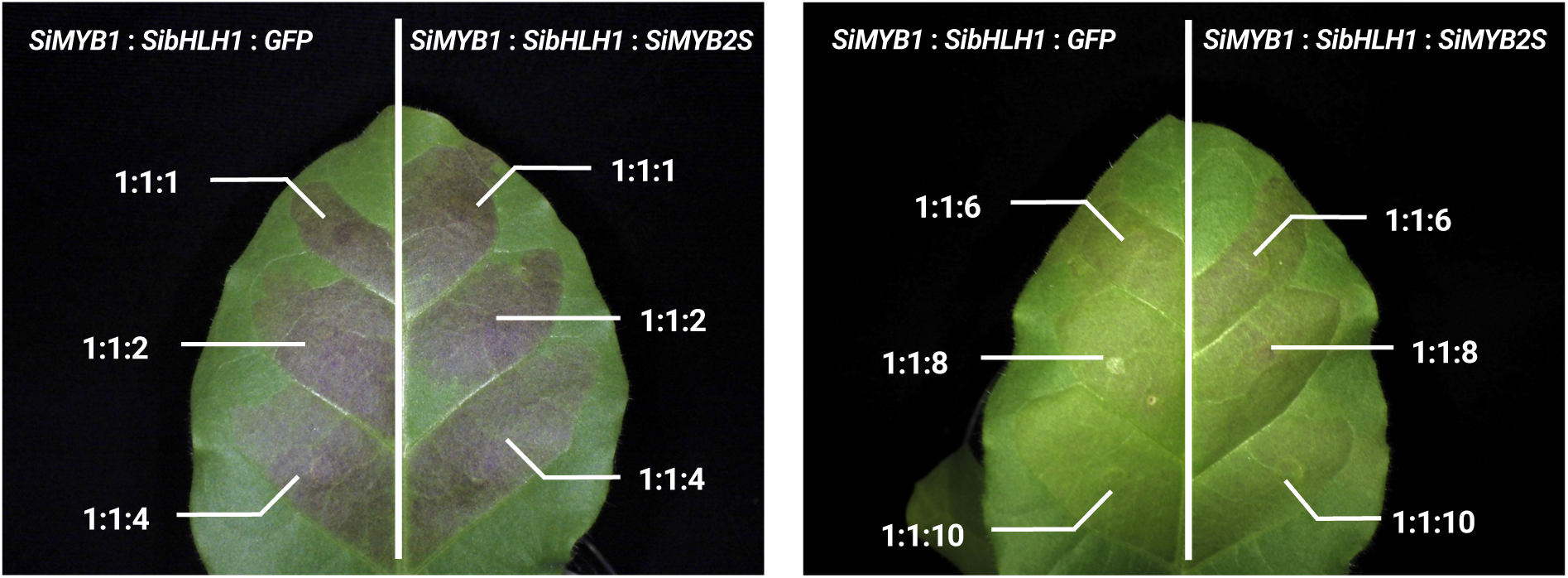
Confirmation of *SiMYB2-Short* repression activity as R3-MYB. The ratio indicates *SiMYB1*: *SibHLH1*: *GFP* (left) or *SiMYB1* (right). Tobacco leaves were photographed four days post infiltration. Infiltration was performed using two leaves from each of the three tobacco plants.

**Fig. S8.**
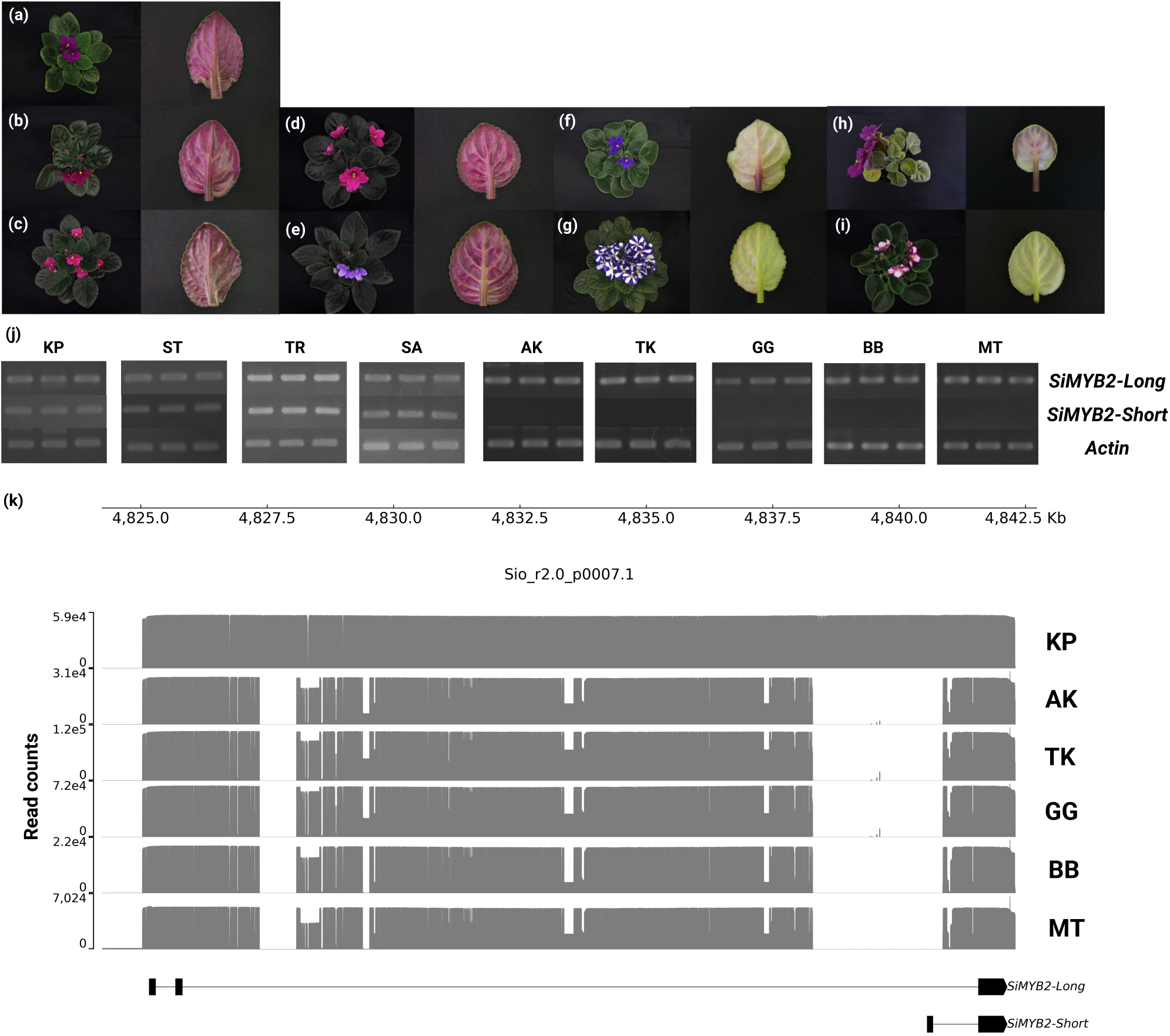
Transcript variants expression and genome structure of the *SiMYB2* gene in several cultivars. (a-i) Rosette and leaf photos of several cultivars used in this study. (a) ‘Akira’ AK, (b) ‘Tomoko’ TK, (c) ‘Georgia’ GG, (d) ‘Barbara’ BB, (e) ‘Manitoba’ MT, (f) ‘Taro’ TR, (g) TR with white-striped petals, (h) ‘Saturn’ SA with red petals, (i) SA. (j) RT-PCR results of *SiMYB2* variants in several cultivars. The amplification reaction was carried out for 40 cycles (plateau). Actin was used as control. (k) Amplicon-seq results in several cultivars. The vertical axis indicates read counts and the horizontal axis indicates the position in the Sio_r2.0_p0007.1 contig.

## Notes

### Summary of Updates

Title was updated; A supplementary table was added to the results as Table S3; Table S3 in the previous version was changed to Table S4.

